# Universal rhythmic architecture uncovers distinct modes of neural dynamics

**DOI:** 10.1101/2024.12.05.627113

**Authors:** Golan Karvat, Maité Crespo-García, Gal Vishne, Michael C Anderson, Ayelet N Landau

## Abstract

Understanding the organizing principles of brain activity can advance neurotechnology and medical diagnosis. Traditionally, brain activity has been viewed as consisting electrical field potentials oscillating at different frequency bands. However, emerging evidence suggests these oscillations can manifest as transient bursts rather than sustained rhythms. Here, we examine the hypothesis that rhythmicity (sustained vs. bursty) adds an additional dimension to brain organization. We segment neurophysiological spectra from 859 participants encompassing a dozen datasets across multiple species, recording techniques, ages 18-88, sexes, brain regions, and cognitive states in health and disease using a novel rhythmicity measure. Combined with simulations and brain stimulation, our results reveal a universal spectral architecture with two categories: high-rhythmicity bands exhibiting sustained oscillations and novel low-rhythmicity bands dominated by brief bursts. This universal architecture reflects stable modes of brain operation: sustained bands suitable for maintaining ongoing activity, and transient bands which can signal responses to change. Rhythmicity thus provides a powerful, replicable, and accessible feature-set for neurotechnology and diagnosis.

## Introduction

Understanding the organizing principles of the brain’s electrical activity using non-invasive techniques is a major goal of neuroscience, with implications for brain-computer interface (BCI) design, pathology diagnosis, and treatment. A dominant hallmark of the electrophysiological signal is its tendency to oscillate. Brain oscillations are thought to signify synchronized fluctuations in excitability, present in neuronal systems from rodents to humans, and can be detected invasively or non-invasively ^1,2^. According to a widely held viewpoint, the entire electrophysiological spectrum is composed of oscillatory bands, covering all frequencies from 0.1 to more than 100 Hz ^3^ [the “canonical bands” delta (0.1-4 Hz), theta (4-8 Hz), alpha (8-13 Hz), beta (13-30 Hz), gamma (30-80 Hz), and high-frequency oscillations (80-250 Hz)]. A century of research and thousands of studies suggest that these seamlessly progressing bands form the spectral architecture of the brain, with oscillatory activity in different bands supporting distinct cognitive and physiological states ^4,5^. This spectral architecture is now considered as one of the fundamental principles of neuronal activity ^6^. However, several practical, functional, and physiological considerations challenge this viewpoint, calling for a revision of this architecture.

On the practical level, band definitions vary widely between research-groups ^7^, which limits our ability to compare findings between studies and species. In addition, bands are currently defined and identified by sustained oscillatory activity and, as such, are assumed to progress seamlessly at the group level. However, at the single-subject level, which is vital for individualized neurotechnology, diagnosis, and treatment, these relatively wide bands consist of a combination of aperiodic activity and some narrow band oscillations ^8^. With no objective framework for relating non oscillatory activity to canonical frequency bands, the role of activity expressed in the gaps between narrow band spectral peaks, if any, remains unknown. On the functional level, at least two of the canonical bands (beta and gamma) are thought to encompass within their range sub-bands with distinct — and even opposing — roles ^9–11^, indicating the need for a more nuanced characterization of bands. On the physiological level, increasing evidence indicates that some oscillatory responses are not sustained. Rather, they arise as strong, short-lived bursts^12–14^.

The bursts raise the possibility that the electrophysiological spectrum segregates into specialised frequency bands according to their persistence, i.e., sustained rhythms versus bursty spectral phenomena in different frequency bands. We therefore hypothesized that neural activity is organized within a rhythmicity-resolved spectral architecture. This architecture is defined not only by frequency but also by a rhythmicity axis that distinguishes between sustained (high-rhythmicity) and bursty (low-rhythmicity) modes of activity. This spectral segregation carries functional significance: it allocates specialized frequency bands to support two distinct processes—the maintenance of ongoing activity, which we hypothesize to be reflected by sustained oscillations in high-rhythmicity bands ^5,15^, and transient responses to inputs, manifested as oscillatory bursts in low-rhythmicity bands ^16,17^.

This rhythmicity-resolved spectral architecture hypothesis yields four testable predictions: (i) There should be defined frequency “sustained bands” that clearly express sustained oscillations. (ii) Additionally, there should be a separate category of bands specializing in transient activity. In these “transient bands”, spectral phenomena should manifest as short-lived bursts. (iii) The architecture is a universal organizing principle of the brain’s operating system; hence it should arise across datasets, recording techniques, brain areas, and species. (iv) if the architecture mediates cognition, its distinct modes of operation should differentially respond to varying cognitive states and input processing.

To test this hypothesis and examine its predictions, we develop an objective standard to divide an individual’s electrophysiological spectrum into high-rhythmic and low-rhythmic bands. We then show that the resulting rhythmicity-resolved spectral architecture is universal and bears functional significance: rhythmic bands signify maintenance of ongoing activity, whereas non-rhythmic bands signify transient events.

## Results

### Rhythmicity-resolved spectral architecture

To test the predictions of the rhythmic architecture hypothesis, we developed a phase-based tool to measure rhythmicity over the frequency spectrum (Fig. 1, fig. S1, and online methods). Three features characterize brain oscillations: frequency (cycles per time-unit), power (or magnitude), and phase (or angle). The canonical bands are typically identified by measuring spectral power at different frequencies (fig. S2A-C). However, estimated power is influenced by both oscillatory activity and non-oscillatory transients. For example, a spectral power peak could result from strong impulses which are not rhythmic. An alternative approach to power is to estimate how sustained the oscillation is by measuring how well, at any moment, the signal’s phase predicts its future phase. This is achieved by computing the coherence, or phase similarity, between the signal and a time-lagged copy of itself (Fig. 1A). Previous authors defined rhythmicity as “the consistency of the phase relations between timepoints that are separated by some interval (lag) ^18^.” Then, they measured the rhythmicity spectrum as a function of lag duration, assuming that genuine oscillations should remain sustained for many cycles ^18–20^ (Fig. 1B). Here we adopt this definition of rhythmicity, and note that the term is not meant to imply the existence of an underlying rhythm-generating circuit, but rather denotes the degree of phase consistency. Based on large datasets and comprehensive simulations, we found that the choice of lag does not influence the relative rhythmicity values obtained across the different frequencies for each individual subject (Fig. 1C). Hence the rhythmicity spectrum can be derived from a single lag.

**Figure 1.**
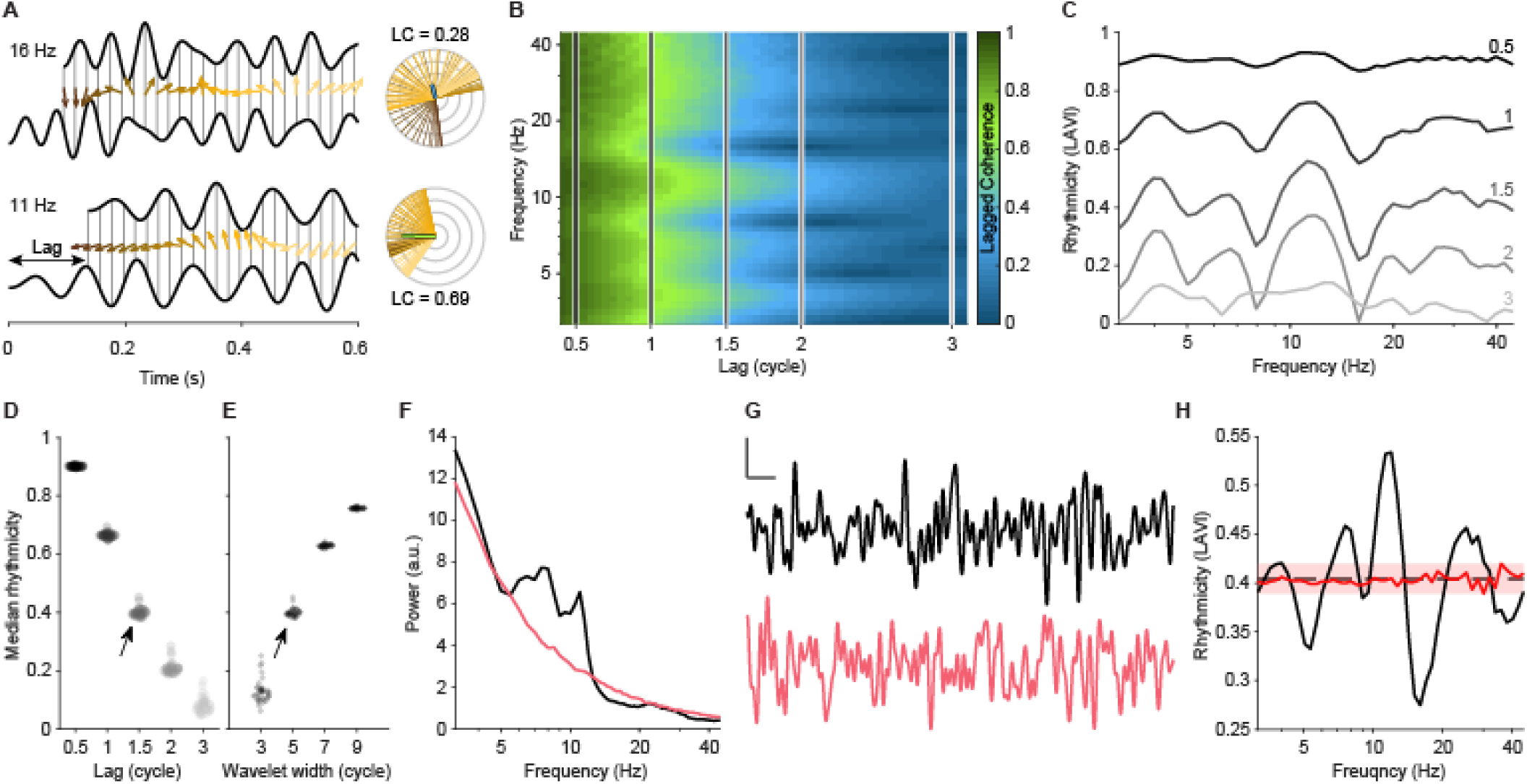
LAVI is a measure of rhythmicity. **A.** To compute the Lagged Coherence (LC) at each frequency, the phase at each timepoint is subtracted from the phase of a lagged copy of the signal (arrows, color-coded by time). LC is the vector mean of phase differences across all timepoints (right, circular histograms). Rhythmic signals yield consistent phase differences and thus high LC values; non-rhythmic signals yield lower LC values. **B.** LC (colour) as a function of lag (abscissa) and frequency (ordinate) of one participant. Vertical lines correspond to specific lags plotted in C. **C.** LC dynamics across lags. Peaks at specific frequencies are found in different lags, with ceiling/floor effects for very short or long lags. To avoid confusion between LC (computed over different lags) and rhythmicity in different frequencies at a specific lag, we term this value Lagged Angle Vector Index (LAVI). **D.** The LAVI median is consistent across participants. Shown are the medians (across frequencies) of each of the N = 37 participants from Dataset I (dots; shades of grey as in C), plotted as a function of lag. For all lags, a wavelet width of 5 cycles was used. Note the relatively low variability across participants at each lag. **E.** As in D, but with varying wavelet widths. Lag was fixed at 1.5 cycles for all participants. Arrows in C and D indicate the parameter values used throughout the manuscript (1.5 cycles lag and 5 cycles wavelet, correspondingly). **F.** Power spectral density of one subject (black) and the aperiodic component estimated using a power-law fit (pink). This fit was used to generate surrogate data with a similar power spectrum but randomized phase structure. **G.** Raw traces from the original (black) and surrogate (pink) data. Vertical scale bar: 10 μV; horizontal scale bar: 0.1 s. **H.** LAVI of real (black) and surrogate (pink) data. Rhythmic peaks and troughs are evident in the real data but absent in the surrogate. Dashed line: median of data. Pink shade: noise ribbon, defined as the range of the surrogate.

We term the algorithm used to resolve rhythmicity across different frequencies at a specific lag the *lagged-angle vector index* (LAVI). LAVI is defined as the vector mean, computed across all time points, of the phase difference between the signal and a lagged copy of itself. As a measure of phase-consistency, it is influenced by the rate of phase-shift occurrence (fig. S1). Importantly, LAVI values show a clear baseline: the median across frequencies is stable across participants, channels, and datasets, and depends only on experimenter-controlled parameters such as lag duration (Fig. 1D) and the window size used for spectral estimation (Fig. 1E). To test whether the LAVI spectrum reflects background noise, we generated surrogate datasets. These were constructed by randomizing phases while matching each dataset’s 1/f power profile—the inverse relationship between power and frequency characteristic of neurophysiological signals (see Methods; Fig. 1F–H). Recorded and surrogate data had similar median rhythmicity values, but the surrogates showed a narrower, flatter range. Leveraging this property, we define the median across frequencies as baseline rhythmicity and the surrogate range as the ‘noise ribbon’ (see Methods and Supplementary Text; fig. S6).

The use of a single lag in LAVI not only accelerates computation by an order of magnitude compared to previous lagged-coherence methods, but also establishes a universal baseline of rhythmicity. This baseline enables the identification of frequency bands exhibiting both increased and decreased rhythmicity, within and across individuals. LAVI is therefore uniquely suited to test our rhythmic-architecture hypothesis by identifying frequency ranges that lie between two crossings of the baseline and above the noise ribbon as high-rhythmicity bands. Conversely, frequencies between two baseline crossings that fall below the noise ribbon are classified as low-rhythmicity bands (Fig. 2A). Integrating LAVI with simulations of surrogate distributions (Fig. 1F-H), we developed an *automated band-border detection algorithm* (ABBA). ABBA delineates the boundaries of significantly high- and low-rhythmicity bands at the individual subject level and automatically assigns band labels. Specifically, we consistently observed a dominant peak within the 6-14 Hz range, consistent with the traditional ‘alpha’ band, and therefore we label this peak is ‘alpha’ (α); the subsequent trough is labelled ‘beta1’ (β1), followed by the next peak as ‘beta2’ (β2), and the following trough as ‘gamma1’ (γ1). Similarly, the trough preceding alpha is labelled as ‘theta/alpha’ (θ/α), the adjacent lower-frequency peak as theta (θ), the next lower trough as ‘delta/theta’ (δ/θ) and the lowest peak considered in this manuscript is labelled delta (δ) (see Fig. 2A).

**Figure 2.**
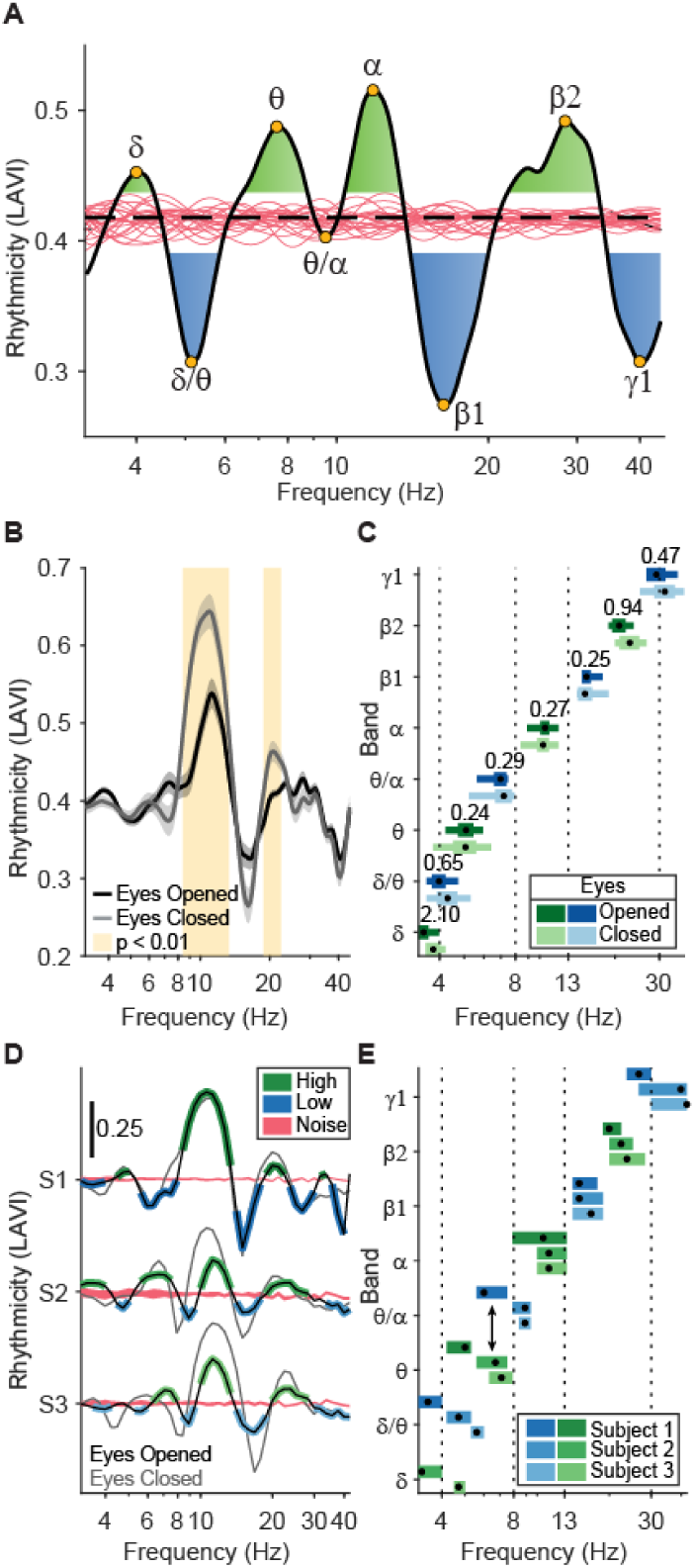
ABBA detects individual and group-level brainwave bands. **A.** Rhythmicity profile of a representative participant, with output of the Automated Band Border Algorithm (ABBA). Black line: subject’s rhythmicity; dashed line: median; pink traces: 20 instances of simulated 1/f noise. Yellow circles mark peaks and troughs. Green and blue areas indicate frequencies with significantly increased or decreased rhythmicity, respectively. **B.** Group-level rhythmicity profiles (N = 37) for eyes open (black, dataset I) and closed (grey, dataset II). Shading: SEM; yellow highlights significant clusters (p < 0.01, cluster-based permutation test). **C.** ABBA-derived group-level band annotations. Dots: mean peak frequency; thin horizontal lines: interquartile range of high- (green) and low-rhythmicity (blue) bands; thick horizontal lines: 95% confidence intervals. Bayes factors (black numbers above each band) quantify group differences between eyes open and closed (BF < 1: no difference). Dashed lines indicate canonical (power-based) bands. Note: While the brain states affect rhythmicity magnitude, the automatically detected band frequencies remain stable. **D.** Rhythmicity profiles from three participants during eyes-open (black) and eyes-closed (grey) conditions. Subject-specific ABBA-identified bands with high (green) or low (blue) rhythmicity during eyes-open session are overlaid. **E.** Automatically detected bands for participants in D. Dots show peak frequencies; horizontal lines span detected band ranges (green: high rhythmicity; blue: low). Note individual variability, including cases where the same frequency is classified differently across subjects (arrow).

At the population level, peak frequencies of high-rhythmicity bands defined by ABBA aligned with canonical, power-based frequency bands (Fig. 2B–C) ^3^. Yet individual subjects showed considerable variability within these ranges, with some frequencies classified as high- or low-rhythmicity depending on the subject (Fig. 2D–E and fig. S2G). This variability underscores the need for an objective, single-subject band-detection framework. Our approach thus enables comparison with canonical band names while capturing the individualized spectral structure that underlies them.

This alignment also enabled us to explore the functional relevance of the spectral architecture and to test our prediction that rhythmicity patterns are modulated by cognitive states. To this end, we examined weather changes commonly observed in oscillatory power across brain states are also reflected in rhythmicity. Specifically, we compared the rhythmicity profiles of the same participants while they were engaged in a visual task and during rest with eyes closed (Fig. 2D–E). Alpha-band power is known to increase during wakeful rest with eyes closed ^21^. Consistent with this, we found that rhythmicity in the alpha-band (8.4–13.3 Hz) differed significantly between these cognitive states (*p*<0.01, cluster-permutation test, Fig. 2D). Moreover, the change in rhythmicity (measured via LAVI) was strongly correlated with the change in alpha power (measured in dB, ρ=0.781, *p*<10^-7^). A significant increase in rhythmicity during eyes-closed was also observed in the beta2 band (18.8-22.4 Hz), but not in beta1 (14.7-17.8 Hz), suggesting distinct functional roles for these sub-bands. Notably, the frequency ranges of bands identified by ABBA, as reflecting high or low rhythmicity, remained stable across both states (*t*_295_ = 1.4, *p* = 0.16, BF_10_ = 0.17), underscoring the robustness of the spectral architecture.

### Low- and high-rhythmicity bands show distinct burst dynamics

Our findings point to a stable spectral architecture defined by frequency and rhythmicity, with distinct functional roles for high-rhythmicity bands. Traditional power-based band-detection approaches often overlook low-rhythmicity bands—but do these bands represent physiologically meaningful processes? Or are they merely trivial by-products of rhythmicity peaks? In order to shed light on this question we turned to sub-second manifestations of the spectral architecture. Specifically, we investigated a putative relationship between rhythmicity and burst dynamics in EEG data.

We defined bursts as power peaks exceeding the 90th percentile (see Methods). Visual inspection in the single-subject level revealed that bursts in high-rhythmicity bands (alpha, beta2) exhibited sustained oscillations (≥±3 cycles) with raw and filtered signals closely matching. Conversely, low-rhythmicity bands (theta/alpha, beta1) dampened within ±1 cycle and exhibited phase differences between raw and filtered signals (Fig. 3A), pointing to a link between shorter bursts and reduced phase-consistency in these bands.

**Figure 3.**
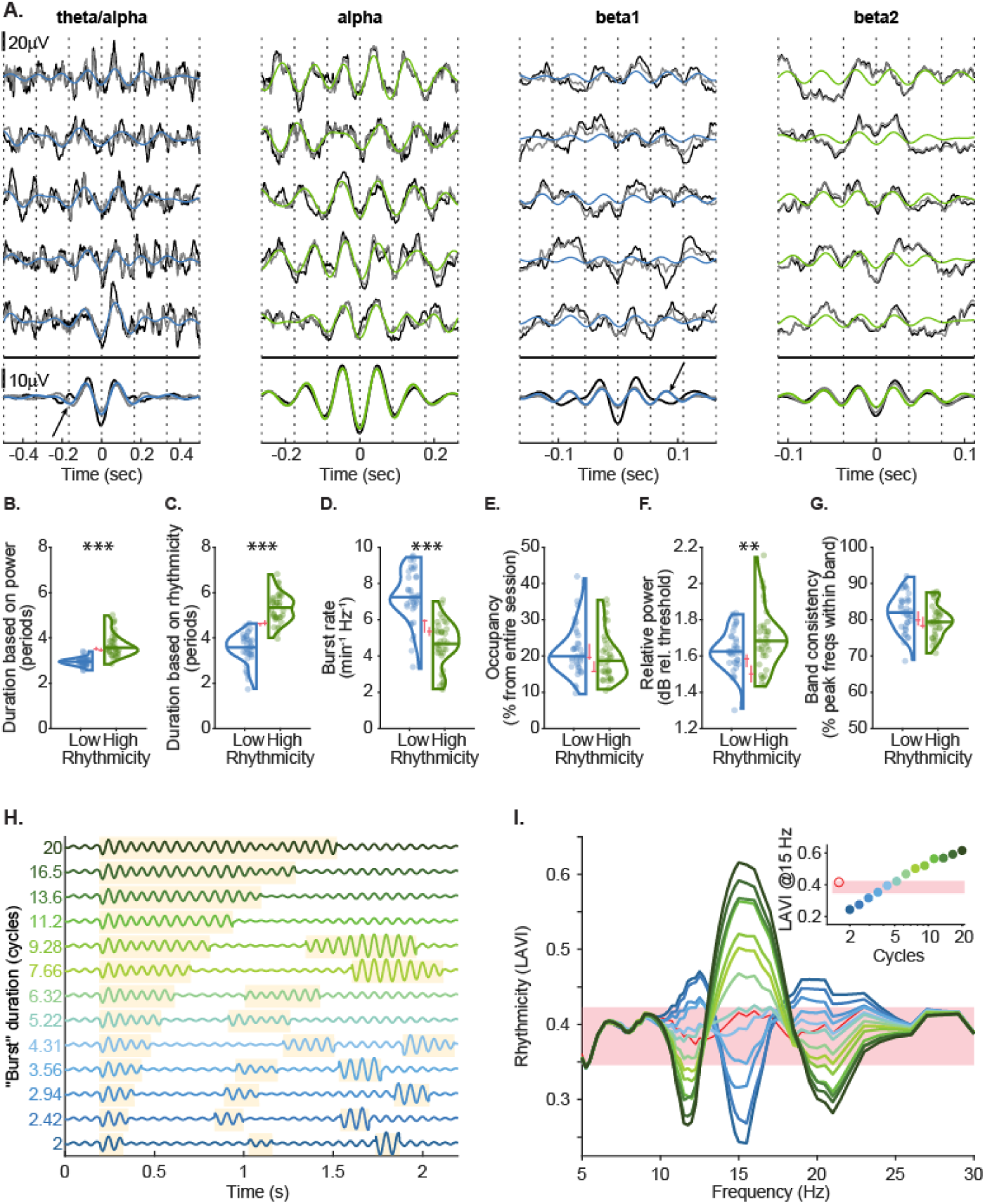
Bursts affect rhythmicity. **A.** Raw bursts in two low-rhythmicity bands (θ/α, β1) and two high-rhythmicity bands (α, β2), aligned to the trough nearest max power (±3 cycles). Black: raw signal (one representative subject). Blue/green: band-pass–filtered. Grey: band-stop filtered (neighbouring bands removed). Bottom row: average across bursts (“burst-related potential”, BRP). Dotted lines mark troughs; arrows indicate phase shifts, likely due to neighbouring high-rhythmicity bands. **B-G**. Burst statistics across low- and high-rhythmicity bands (N = 37): **B**. Duration based on power > 75^th^ percentile. **C**. Duration based on rhythmicity (Within-Trial Phase Lock; WTPL > 75^th^ percentile). **D**. Bursts per minute. **E**. Occupancy (proportion of time in bursts). **F.** Peak-power relative to the 90th percentile threshold. **G**. Frequency overlap (proportion of burst time with peak frequency in-band). Dots: participant means; violins: distributions; horizontal line: medians. Pink element: medians and CIs from 1/f-matched random-phase simulations. **p < 0.01; ***p < 0.001. **H**. Simulated bursts: pink noise was generated from 1/f filtered white-noise. At 15 Hz, filter weight doubled at time-points chosen as bursts (variable length, yellow shading), halved otherwise. **I**. Rhythmicity profiles of simulated data. Trace colour: burst duration (corresponding to H and the inset). Pink shading: LAVI range for unmodulated pink noise; red: unadjusted 1/f noise. Inset: LAVI at 15 Hz.

At the population level (N=37 subjects; see also fig. S3 for analysis across six EEG datasets (total of 178 subjects)), bursts in high-rhythmicity bands lasted significantly longer (3.5 vs. 2.9 cycles), a pattern amplified when using Within-Trial Phase-Lock (WTPL ^22^), a time-resolved measure of phase-consistency (5.3 vs. 3.6 cycles; Fig. 3B-C. See Supplementary table S1 for full statistical details). Conversely, low-rhythmicity bursts occurred more frequently (Fig. 3D), resulting in comparable total durations during which signals exceeded burst thresholds across bands (Fig. 3E), with high-rhythmicity bursts showing slightly higher normalized amplitudes (Fig. 3F). Moreover, although instantaneous frequencies can drift within bursts ^17,23^ (fig. S4), in both modes over 80% of peak frequencies during bursts were confined within the LAVI-defined frequency band (Fig. 3G).

To rule out the possibility that these effects arose from 1/f (“pink”) noise, we compared the EEG data to phase-randomized surrogate signals with matched 1/f power profiles. The simulated signals showed no differences in burst rate or duration across frequencies (Fig. 3B–D, pink lines), and critically, the recorded bursts differed significantly from these noise simulations. In a separate set of simulations we generated bursts with varying length and quantified the resulting rhythmicity of the signal. Bursts shorter than four cycles yielded rhythmicity below chance levels, while bursts longer than six cycles exceeded it (Fig. 3H– I and fig. S4), supporting a distinction between transient and sustained rhythmicity. Together, these findings suggest that bursts are epochs of *band-confined* activity: low-rhythmicity bands reflect brief, transient bursts, whereas high-rhythmicity bands exhibit fewer but longer-lasting oscillations. This raises the question: does this rhythmic architecture hold universally across contexts and recording modalities?

### The rhythmic architecture is universal

Establishing the rhythmicity-resolved spectral architecture as a universal phenomenon requires a robust, diverse, and large sample. Therefore, we computed the rhythmicity spectra of data from 12 datasets acquired from 2450 recording sites of 11 rats and 848 humans (with both invasive and non-invasive methods; Table 1). Using LAVI and ABBA, we found the rhythmicity architecture consistently in all datasets, and across ages, sexes, species, brain regions, and recording techniques, in health and disease.

**Table 1.** Details of datasets used in this study.

| Dataset | Task/ population | Recording technique | N subjects | Recording sites | Age mean (range) | Ref. |
| --- | --- | --- | --- | --- | --- | --- |
| I | Visual duration judgement | EEG | 37 volunteers | Cz | 22.9 (18-29) | 22 |
| II | Eyes closed | EEG | 37 volunteers | Cz | 22.9 (18-29) | 22 |
| III | Tactile duration judgement | EEG | 29 volunteers | Cz | 24 (19-31) | 24 |
| IV | Audio–visual illusion | EEG | 24 volunteers | Cz | (18-42) | 25 |
| V | Memory-control | EEG | 27 volunteers | Cz | 25 (20-34) | 26 |
| VI | Memory-control | HD-EEG | 24 volunteers | Cz | (18-35) | 27 |
| VII | TMS | EEG | 6 volunteers | F4 | 27 (19-37) | New |
| VIII | Rest | MEG | 625 volunteers | 4 magnetometers surrounding Cz | (18-88) | 28 |
| IX | Epilepsy patients | ECoG | 10 patients | 867 electrodes (sites: Fig. 1I) | 41 (19-65) | 29 |
| X | Epilepsy patients | EEG + SEEG | 12 patients | 733 electrodes (sites: Fig. 1J) | 27.5 (17-39) | 30 |
| XI | PD patients | LFP | 17 patients | STNs from 30 hemispheres | 58 (48-65) | 31 |
| XII | Rats navigation | LFP | 11 rats | Hippocampus |  | 32 |

EEG: electroencephalogram, HD-EEG: high-definition (high-channel-count) EEG, TMS: transcranial magnetic stimulation, MEG: magnetoencephalogram, ECoG: electrocorticography, SEEG: stereoelectroencephalogram (depth electrodes), LFP: local field potentials (depth electrodes), Cz: midline central electrode, F4: right frontal electrode, Ref.: reference. PD: Parkinson’s disease.

To determine whether the rhythmic architecture generalizes across recording modalities and brain regions, we analysed spectral profiles from a large dataset comprising both invasive (ECoG, SEEG) and non-invasive (EEG) recordings. We compared the distributions of peak frequencies across datasets (EEG, Fig. 4A–C) and neuroanatomical locations (ECoG, Fig. 4D–F; SEEG, Fig. 4G–I). Using ABBA, we consistently detected the canonical sustained bands—θ, α, and β2—as well as the transient bands—θ/α, β1, and γ1—in nearly all participants (Fig. 4B, 4E, 4H). Detection rates were lower for δ and δ/θ, likely due to our choice of 3 Hz as the lowest frequency analysed, which might not capture δ and δ/θ rhythmicity, resulting in a noisy estimate at these frequencies (i.e., the actual peaks may be in lower frequencies than measured). Importantly, for most participants and electrodes, at least 6 out of 8 bands were detected (Fig. 4C, 4F, 4I). To assess the statistical significance of these bands efficiently, and based on characterization of the parameters affecting the noise ribbon (fig. S6), we first generated 200 random-phase surrogate datasets to construct a reference table of null distributions, matched to a range of 1/f power profiles and acquisition parameters. For each participant and electrode, we then selected the appropriate thresholds from this table based on their individual power spectra, session duration and sampling-frequency. These thresholds were applied within a Monte Carlo framework to determine which observed bands exceeded chance levels. Across the 3–45 Hz range and at α = 0.05, the vast majority of detected bands were found to be statistically significant. Notably, while differences in peak frequencies between datasets and brain regions were observed within bands, the boundaries between bands remained stable—indicating that the rhythmic architecture is robust.

**Figure 4.**
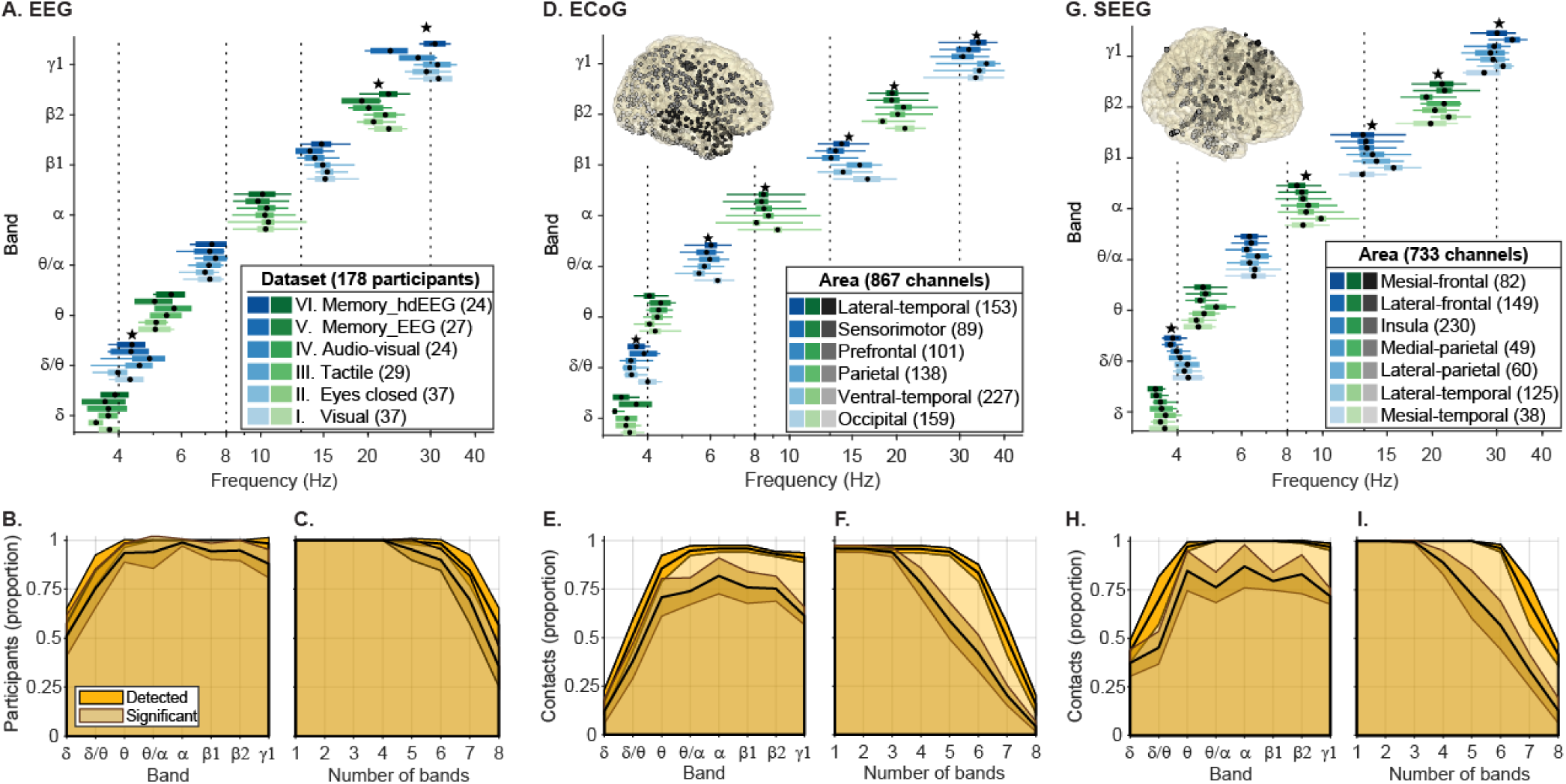
The rhythmicity architecture is universal. **A–C.** Peaks and borders of rhythmicity bands across six EEG datasets (N = 178). **A.** Mean peak frequencies (dots) of high- (green) and low-rhythmicity (blue) bands; thick horizontal bars: 95% CI of the peaks; thin horizontal bars span from the mean of the lowest to the mean of the highest frequency in each band; stars: significant differences across datasets (ANOVA, p < 0.05). **B.** Proportion of participants with each band detected (brown) and statistically significant (yellow); lines: mean; shading: ±SD across studies. **C.** Cumulative proportion of participants with ≥x detected bands; colour as in B. **D–F.** Same metrics for 867 ECoG contacts across six brain regions (dataset IX; legend in D shows region names and electrode counts; SD in E–F over regions). **G–I.** Same for 733 SEEG contacts across seven regions (dataset X).

These consistent patterns across modalities and regions suggest that the rhythmic architecture may help reduce uncertainties arising from methodological and species-related differences. For instance, whether frequency bands identified in invasive recordings such as ECoG or SEEG correspond directly to those observed in non-invasive EEG remains an open question, due to fundamental differences in spatial resolution, depth sensitivity, and susceptibility to signal distortions ^1,33^. To address this, we leveraged a unique recording setting including both scalp EEG and invasive electrodes in dataset X, allowing us to directly compare rhythmicity profiles and individually defined bands between measurement modalities. We found that while the average frequency profile of scalp electrodes tended to be higher than the invasive counterpart (i.e., slightly faster; Fig. 5A) relative to invasive electrodes, the individual band peak frequencies identified using ABBA were remarkably similar across measurement modalities (BF_10_=0.042; BF_01_=23.8), providing strong evidence in favor of the null hypothesis of no difference between modalities (Fig. 5B).

**Figure 5.**
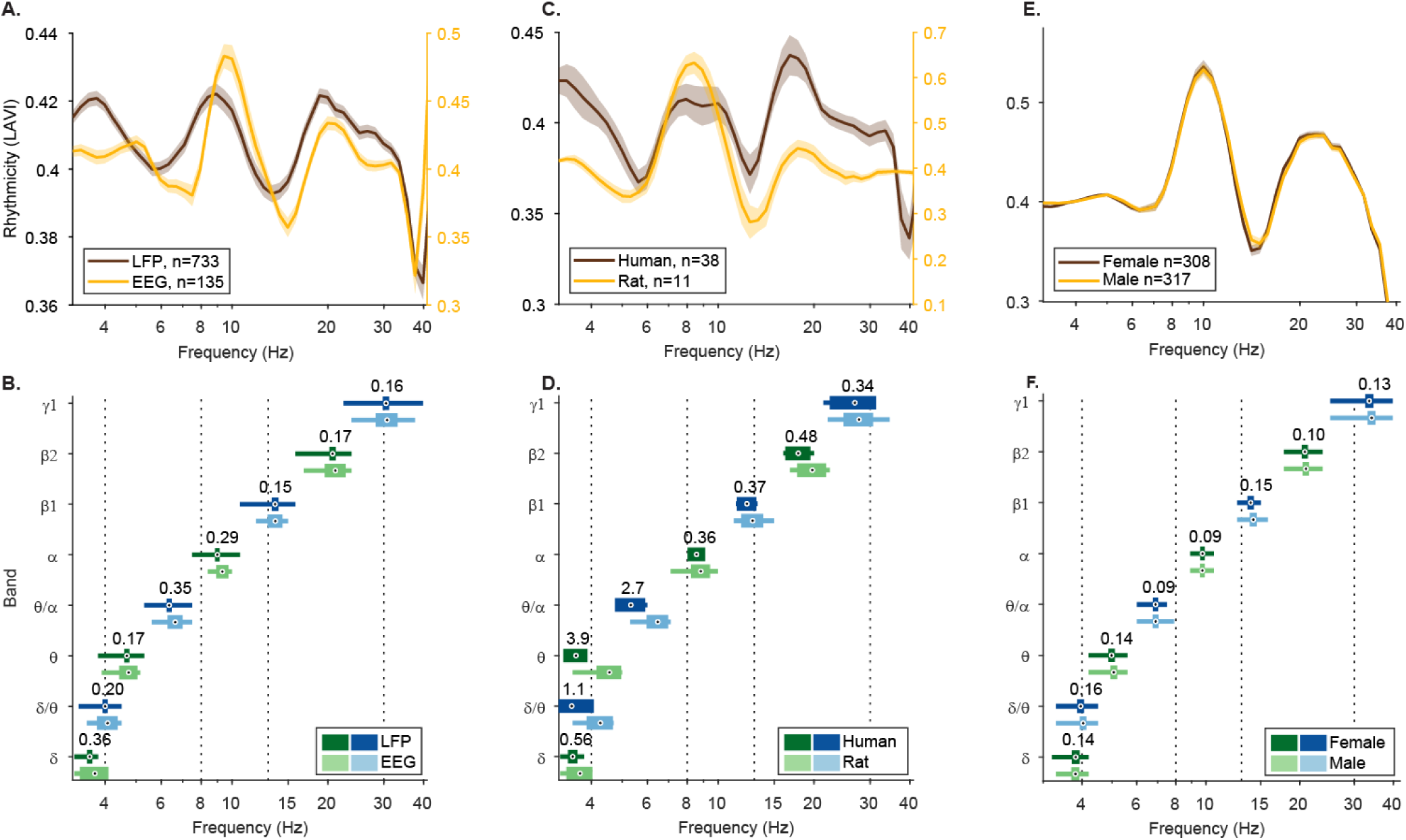
Rhythmicity reduces uncertainties in existing data. **A, B**. LAVI (A) and ABBA (B) analyses of concurrently recorded LFP and EEG in patients (dataset X) reveal that population averaged EEG band peaks tend to be faster (rightward shift of yellow vs. brown line). However, at the individual level, Bayes factor values below 1 indicate similarity between recording methods. **C-D**. Comparison between human hippocampal SEEG (dataset X) and rat hippocampal LFP (dataset XII). Bayes factor analysis provides evidence for no difference between species in Alpha, Beta1, Beta2, and Gamma1 bands **E-F**. The rhythmicity architecture of female and male participants from dataset VIII is similar. In A, C, and E lines indicate means and shaded areas represent SEM; numbers in denote electrode counts. In (B, D, F), dots mark mean peak frequencies per band, with thick horizontal bars showing 95% confidence intervals and thin bars the interquartile range. Black numbers above each band show between-group Bayes factors from t-tests.

Furthermore, the invasive recordings from patients’ hippocampi enabled a cross-species comparison of rhythmic architecture with rats. A longstanding debate concerns whether human alpha oscillations (8–12 Hz ^3^) are functionally analogous to rodent theta (4–12 Hz ^34^). While some argue these rhythms reflect similar mechanisms despite frequency differences ^35,36^, others question whether rodents exhibit alpha at all, noting their hippocampal activity is dominated by theta ^37,38^. To address this and test whether the rhythmicity architecture generalizes across species, we analyzed hippocampal recordings from 11 rats (Dataset XII). As in humans, both sustained and transient bands from θ to γ1 were detected in nearly all rats (100% for θ–γ1; significant in 100% for θ/α, α, and β1; 90% for β2; 82% for θ and γ1). Although rhythmicity magnitudes were higher in rats (note scale in Fig. 5C), band frequency distributions were strongly conserved across species (BF_01_ = 10.99, Fig. 5D). Notably, the dominant peak in rats was around 8 Hz—labelled ‘theta’ in the original publication of dataset XII ^32^, but identified as ‘alpha’ by ABBA— suggesting that rodent theta reported in the literature may correspond to human alpha, as the most prominent rhythmic band below ∼15 Hz. To further assess the generalizability of rhythmicity architecture, we examined potential sex differences in Dataset VIII. LAVI and ABBA analyses (Fig. 5E–F) showed comparable rhythmicity profiles and band peak frequencies between female and male participants, with strong evidence for no difference (BF_01_ = 43.25).

Taken together, these findings reveal a dual-mode rhythmicity architecture: sustained bands characterized by long-lasting oscillations, and transient bands dominated by brief bursts. This architecture generalizes across datasets, brain regions, recording modalities, sexes, and species, supporting the idea of a universal organizing principle. We now turn to our final prediction: that these distinct rhythmic modes should differentially reflect and respond to varying brain states and input processing demands.

### The rhythmic architecture reflects brain states

To explore how rhythmic modes reflect brain state, we first investigated age-related changes in rhythmicity magnitude and peak frequency. Aging is associated with widespread changes in brain structure and function, many of which are linked to cognitive decline ^39^. Neural oscillations, particularly in the alpha band (∼8–12 Hz), are known to decelerate and reduce power with age, correlating with reduced processing speed and diminished attentional control ^40^. However, more recent studies suggest that such effects may partly stem from changes in the aperiodic background signal ^41^. Aging thus provides a valuable testbed for examining whether the rhythmicity architecture reflects changes in brain state. Because LAVI and ABBA are less influenced by variations in the 1/f slope—owing to their use of a robust baseline— they enable assessment of age-related changes without confounds that have limited previous analyses ^41^. We analysed rhythmicity profiles from magnetoencephalography (MEG) recordings of a large cohort of healthy individuals spanning the adult lifespan (N = 625, ages 18-88; Dataset VIII ^28^). We observed age-related reduction in rhythmicity magnitude and frequency not only in alpha but also in other sustained (delta, theta, and beta2) and transient bands (theta/alpha and beta1; Pearson correlation per participant and ANOVA per age decade group, Fig. 6A and 6B and Table S1). These findings indicate that both sustained and transient rhythmic modes are sensitive to age-related brain changes, underscoring their utility as markers of brain state across the lifespan.

**Figure 6.**
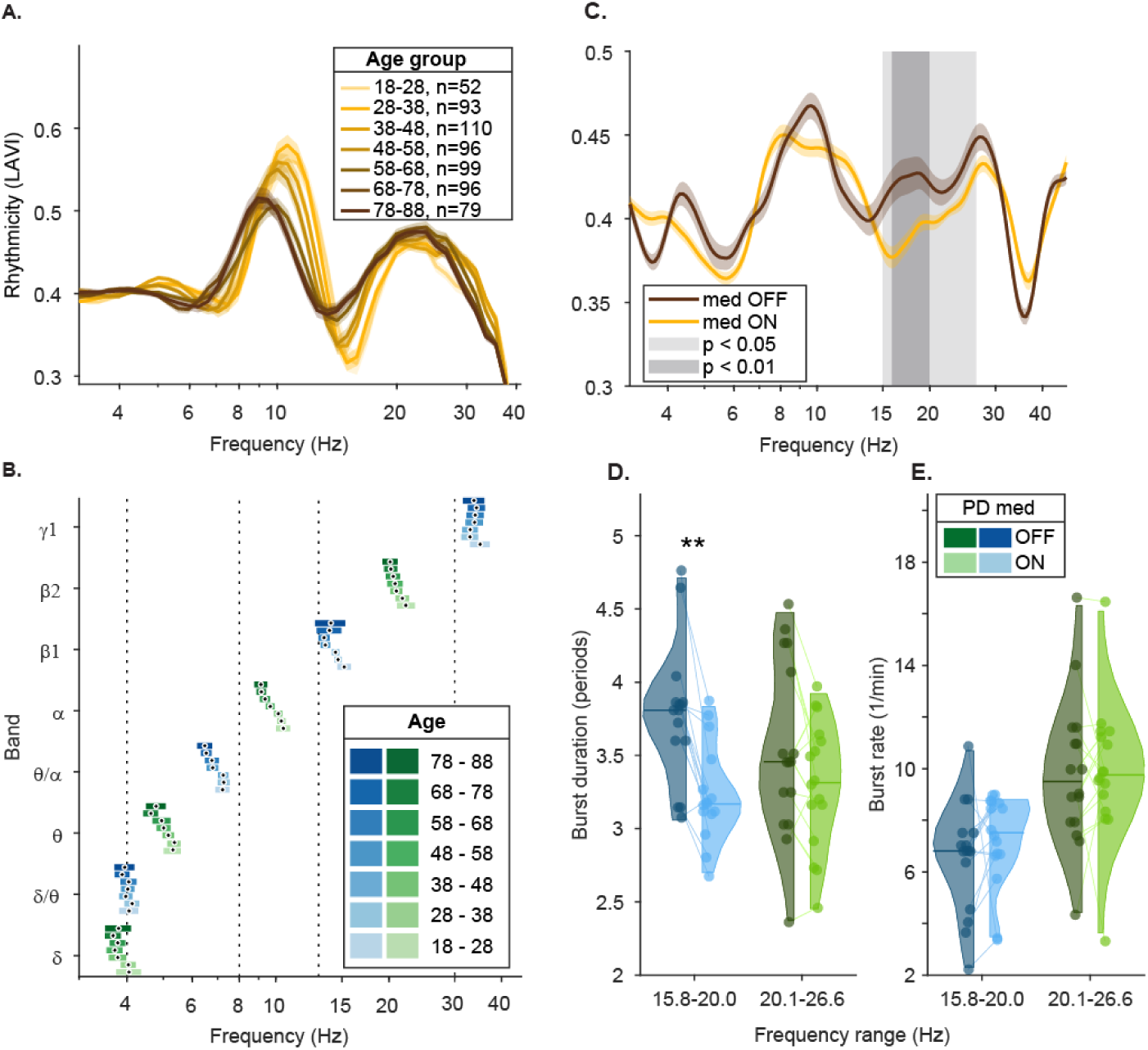
Rhythmicity changes in healthy-aging and disease. **A, B.** LAVI (A) and ABBA (B) analyses over the adult lifespan (Dataset VIII). In A, lines indicate group means and shaded areas represent SEM; numbers denote participant count per age group (in years). In B, dots represent mean peak frequencies, horizontal bars: 95% confidence intervals. **C.** Population-averaged rhythmicity profiles from Sub-thalamic Nuclei (STN) of PD patients (N = 17; 30 hemispheres, Dataset XI), comparing ON vs. OFF L-Dopa states. Grey shading: significant clusters from permutation analysis. **D, E.** Burst duration (D), defined as power > 75^th^ percentile, and number of bursts per minute (E) from hemispheres showing increased beta1 rhythmicity OFF vs. ON medication (15.8–20.0 Hz, median split). Dots represent individual STN means; violins show distributions; horizontal bars denote medians. *p < 0.05; **p < 0.01; ***p < 0.001, Pearson correlation with false discovery rate according to the Benjamini-Hochberg algorithm (B) or ANOVA with Fisher’s least significant difference (LSD) post-hoc (D).

This large-scale group characterization provides a normative reference for neuro-typical aging. We next asked whether the rhythmic architecture is sensitive also to pathological brain states. To address this, we compared rhythmicity profiles of LFP recordings from sub-thalamic nuclei (STN) of Parkinson’s disease (PD) patients, with (i.e., “ON”) and without (i.e., “OFF”) medication (N = 17 patients, 30 hemispheres; Dataset XI). In PD, beta power (∼13-30 Hz) is known to increase ^42^. Activity in the beta band is believed to be controlled by the level of dopaminergic activity in response to internal and external cues, and following dopaminergic cells’ loss, beta levels are elevated in PD ^43^. We found a significant increase in rhythmicity OFF medication compared to ON in the entire beta-band (15.0 - 26.6 Hz, *p*<0.05, cluster permutation test; Fig. 6C). Notably, using a permutation test with stricter threshold of p<0.01 the medication effect was localized to the transient band beta1 (15.8 – 20.0 Hz, *p*<0.01). This suggests that while PD pathology elevates rhythmicity across the beta range, the effect is particularly pronounced in the transient band beta1, implicating burst dynamics in the pathophysiology. Indeed, prior studies have linked increased STN beta power to prolonged beta bursts, which are shortened in response to medication ^44,45^. To further explore whether the pathological burst dynamics are related to rhythmicity modes, we compared burst duration and rate in beta1 (15.8 – 20.0 Hz) and beta2 (20.1 - 26.6 Hz) from STNs exhibiting medication-related beta1 power modulation (median split). We found significantly longer burst durations OFF medication specifically in beta1 (*p* < 0.01; Fig. 6D). No significant differences were observed in burst durations for beta2 or in burst rates for either beta band (Fig. 6E; all *p* > 0.15).

### Dual Rhythmic Modes and Input Processing

The modulation of rhythmicity by engagement, age, and disease (Figs. 2D and 6) suggests a recurring pattern of sensitivity to brain states. To investigate how this architecture supports cognition, we next examined how its distinct modes operate on shorter timescales during input processing. Specifically, we asked whether sustained rhythms, being stable over many cycles, support ongoing maintenance, while transient rhythms, being brief and event-linked, reflect responsiveness to change. To test this, we employed two complementary approaches: responses to visual stimuli recorded from invasive ECoG electrodes, and responses to transient, rhythmic, and arrhythmic TMS stimuli recorded using EEG.

Participants in Dataset IX (N = 10) performed a visual perception task while neuronal activity was recorded using electrocorticography (ECoG). As previously reported ^29^, analysis of high-frequency (HF; 70–150 Hz) signal amplitude—an established proxy for neuronal firing—revealed that among 157 contacts over the occipital cortex, 90 (57%) exhibited increased HF activity in response to a brief (300 ms) visual stimulus, while 36 (23%) showed decreased activity (Fig. 7A; 31 contacts [20%] were non-responsive and are not shown). We consider the increased-HF channels as positively responding to the input and engaging in its processing and the decreased-HF channels as negatively responding, potentially engaged in other, ongoing processes. Thus, according to the framework of the rhythmic architecture, we predicted the former group to be coupled with transient bands, and the latter with sustained bands. Furthermore, in the absence of input, we expect neuronal firing of both groups to correlate with activity in sustained bands.

**Figure 7.**
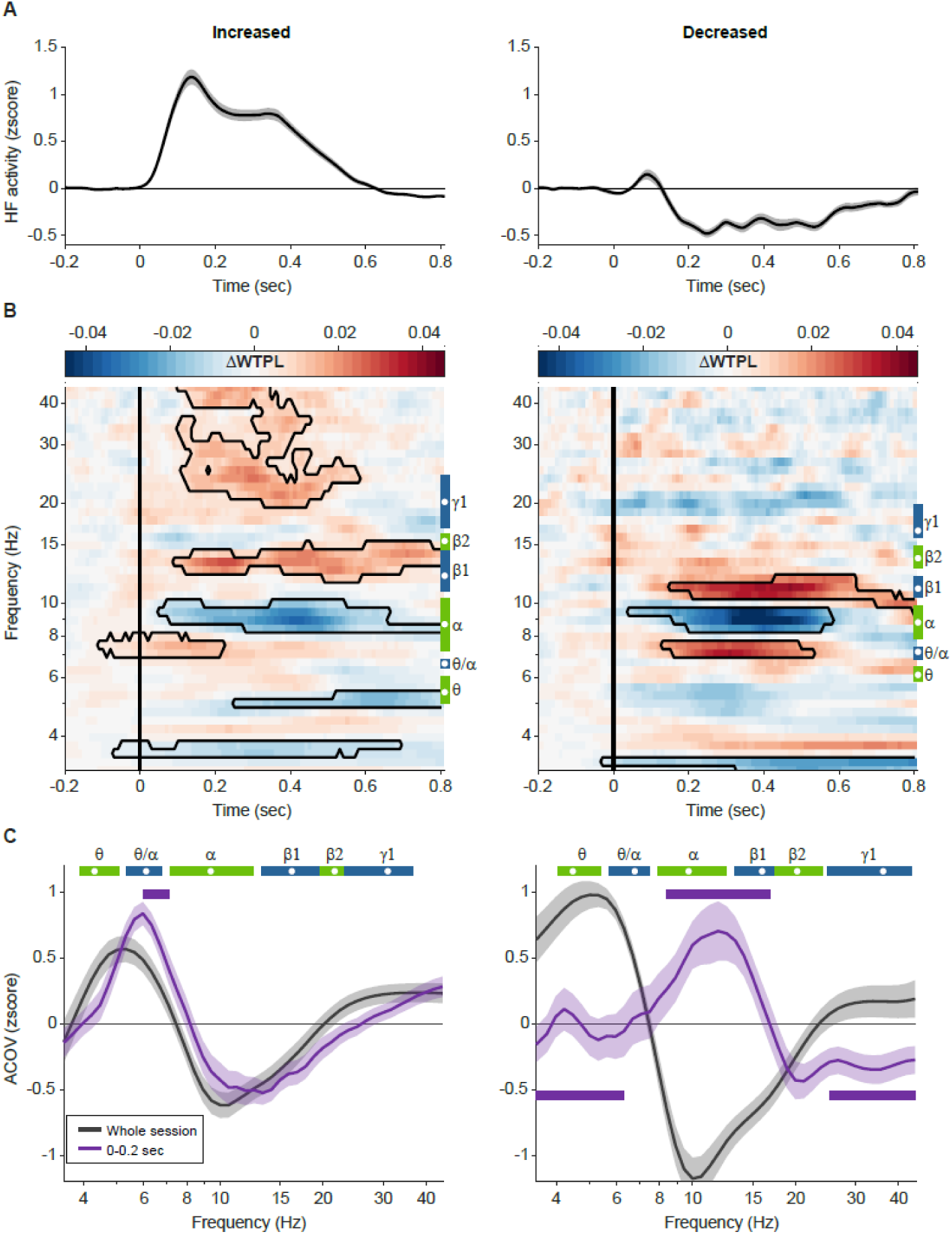
Band-specific responses to visual stimulation. **A.** Visual stimulus presentation elicited changes in high-frequency activity (HF; 70–150 Hz, a proxy for firing rate) over occipital ECoG contacts (Dataset IX). Of these, 57% (n = 90) showed an HF increase (left), while 23% (n = 36) showed a decrease (right). Panels B-C are also divided to increased (left) and decreased (right) units. **B.** Within-trial phase-locking values (WTPL) across frequencies, normalized to a 0.3 s pre-stimulus baseline. Vertical blue and green vertical element on the right side of each panel indicate mean frequencies of transient and sustained bands across participants, respectively, computed as the WTPL mean over time during the baseline; white dots mark the mean peak frequency per band. Black contours denote clusters with p < 0.01 (cluster-based permutation test on t-values). Notably, ΔWTPL increased in transient bands (θ/α, β1) and decreased in sustained bands (θ, α). **C.** Low-frequency (LF)–HF amplitude covariance (ACOV). Grey: ACOV computed over the whole session; purple: ACOV during 0–0.2 s post-stimulus. Horizontal lines indicate significant differences (p < 0.01) between the stimulus period and the session baseline. Band marks (horizontal blue and green elements at the top of the panels) were computed using ABBA over the whole session (similar to B).

To explore how these two neuronal subpopulations relate to rhythmicity in response to input, we computed the average within-trial phase-locking value (WTPL) aligned to stimulus presentation (Fig. 7B). As previously reported ^22^, sustained-bands exhibit high WTPL values at baseline. We leveraged this property to compute the task-specific rhythmic profile as the mean WTPL during the 300 ms preceding stimulus onset. We then assessed how rhythmicity changed in response to input by computing the WTPL normalized via baseline subtraction (ΔWTPL). In both channel types (increased and decreased HF), we observed increased rhythmicity in transient bands and decreased rhythmicity in sustained bands following stimulus onset. The two subpopulations diverged at frequencies above ∼20 Hz, where channels with increased HF activity exhibited elevated rhythmicity.

Next, we sought to investigate how neuronal firing-rate and ongoing oscillatory dynamics relate to cognitive states. Here too, we use the HF as a proxy for firing-rates and quantify the covariance between HF amplitude and the amplitudes of each of the lower frequencies (LF, 3-45 Hz). We predicted that positively responding contacts would exhibit stronger HF-LF covariation within transient bands upon stimulus presentation, reflecting their participation in input processing. In contrast, contacts that are inhibited by the stimulus would exhibit stronger HF-LF covariation within sustained oscillation bands, reflecting their involvement in computations other than the transient input. We found that over the whole session, where in general the amount of input is low, HF significantly covaried more with activity in the sustained band theta (defined by LAVI and ABBA; Fig. 7C, grey traces; increased-HF channels: 5.0-6.0 Hz; decreased-HF: 5.8-6.4 Hz), and less in alpha (increased-HF: 7.2-10.3 Hz; decreased-HF: 7.8-9.8 Hz). In contrast, when focusing on the time of visual input (0 to 0.2 s after stimulus onset; Fig. 7C, purple traces), positively responding channels increased their HF-LF covariance at the transient band theta/alpha (Fig. 7C, left panel; frequencies significantly [*p* < 0.01] higher than whole session: 5.9-7.1 Hz). Negatively responding channels, conversely, shifted their strongest HF-LF covariance into the alpha range (with additional effects in beta1: 8.4-16.8 Hz; Fig. 7C, right panel), supporting a role of alpha in inhibition ^46,47^.

Having observed how transient discrete inputs are associated with rhythmicity, we next examined how repetitive inputs—both rhythmic and arrhythmic—relate to the architecture. To this end, we used Transcranial Magnetic Stimulation (TMS) to deliver transient, rhythmic, and arrhythmic inputs (Fig. 8A). Single TMS pulses evoked characteristic EEG responses consisting of a sequence of peaks and troughs (“ripples”; Fig. 8B). Consistent with previous studies ^48^, this response was reflected in increases in the power across a wide range of frequencies (Fig. 8C). However, according to the rhythmic architecture framework, the rhythmic response to brief events should be confined to the transient bands. To test this, we measured time-resolved rhythmicity (WTPL), and found that ripples were associated with significantly increased rhythmicity specifically in the transient bands θ/α, β1, and γ1 (*p*<0.05, cluster permutation test; Fig. 8D), lasting up to 300 ms post-stimulation.

**Figure 8.**
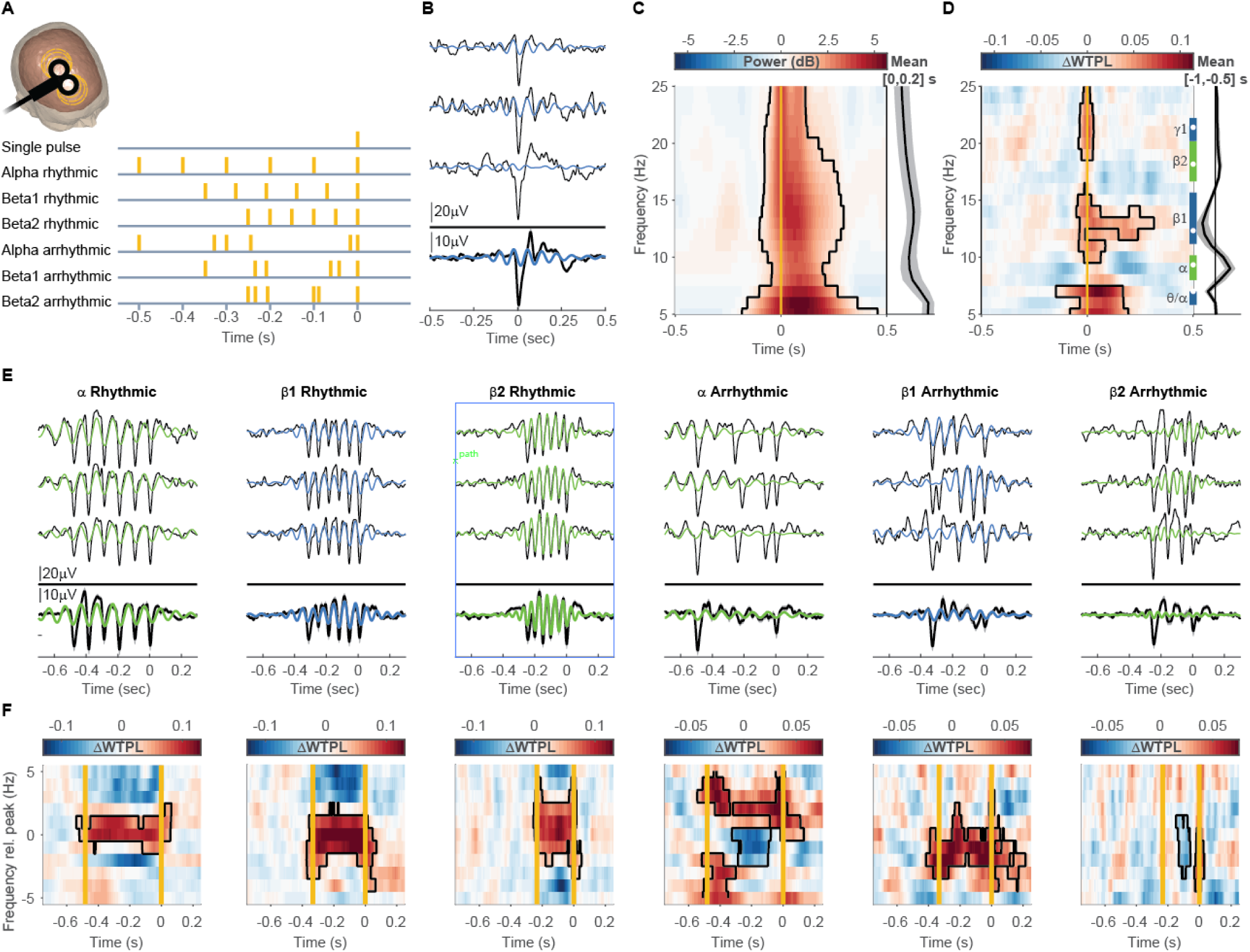
Band-specific responses to TMS stimulation. **A.** Stimulation protocol. TMS over the right dorsolateral prefrontal cortex (rDLPFC) alternated randomly between single pulses and trains of six rhythmic or arrhythmic pulses (yellow vertical lines). **B.** Single-trial responses (top three traces) and the averaged response (bottom trace) to single-pulse stimulation from a representative participant. Black: raw signal; blue: signal filtered in the beta1 band. Shaded area represents ±SEM (barely visible due to low variation levels). **C-D.** Population averaged (N = 6) power (C) and Within-Trial Phase-Locking value (WTPL; D) at each frequency, normalized to a 0.5–1 s pre stimulation baseline. Trace plotted to the right of the panel depicts the mean power during the first 200 ms after TMS (C) or baseline rhythmicity during the [−1 −0.5] s baseline window (D). Blue and green vertical elements indicate the mean frequencies of transient and sustained bands across participants, respectively. White dots mark the mean peak frequency per band. Black contour denotes clusters with p < 0.05 (cluster-based permutation test over t-values). Yellow line at time = 0: TMS single-pulse. Notably, following the TMS pulse, power increases in a wide range of frequencies, while WTPL values increase specifically at transient bands. **E-F.** Repetitive TMS. E. Single-trial responses (top three traces) and the averaged response (bottom trace). Black traces: raw signal; coloured time-series (green and blue) indicate signals filtered at the band used for rTMS. F. WTPL. Yellow vertical lines mark the first and last TMS pulses. Frequencies (ordinate) are shown relative to the individual peak frequency of each participant in each band. Black contour: significant (*p* < 0.05) cluster, permutation test.

For repetitive inputs, we predicted that rhythmic stimulation (i.e. fixed inter-pulse interval [IPI]) would increase EEG-measured rhythmicity, whereas arrhythmic stimulation (random IPI) might violate phase consistency and thus disrupt ongoing oscillations, particularly in sustained bands. To test this, we computed each participant’s rhythmicity profile at rest and determined their individual alpha, beta1, and beta2 peak frequencies. For each band, we then delivered 60 rhythmic trains of six pulses each, with IPIs fixed to match the cycle length of the band’s peak frequency. The arrhythmic condition also consisted of 60 trains of six pulses, matched in total duration between the first and last pulses but with random temporal placement of the pulses within the sequence. Similar to single pulses, each repetitive impulse induced a strong peak and valley (Fig. 8E). In the rhythmic condition, these individual responses were temporally aligned with the band’s oscillations (note phase-lock between pulses and the filtered EEG signal in Fig. 8E). As a result, WTPL values during rhythmic stimulation significantly increased relative to pre-stimulus baseline in all bands (Fig. 8F).

Arrhythmic repetitive stimulation induced band-specific effects. In the sustained bands alpha and beta2, arrhythmic stimulation decreased rhythmicity relative to baseline (blue clusters around peak frequency (0), Fig. 8F), implying that the arrhythmic pulses disrupted ongoing oscillations. In beta1, however, arrhythmic stimulation increased rhythmicity relative to baseline up to 200 ms after the last pulse. This likely reflects the summed contributions of individual pulse responses, as each pulse responses contains a prominent beta1 component (see single-pulse results in Fig. 8D). Together, these results support our hypothesis that low-rhythmicity bands consist of transient yet spectrally defined activity driven by discrete inputs, whereas high-rhythmicity bands reflect sustained oscillations.

## Discussion

In this study we uncover a universal rhythmic architecture composed of two types of bands – sustained and bursty – which alternate intermittently along a logarithmically scaled frequency axis. Rhythmicity (i.e., sustained and bursty) is revealed as a fundamental and previously overlooked dimension of brain dynamics (Fig. 9). This functional segmentation of spectral activity provides a principled basis for defining frequency bands, which is essential for aligning methodologies across studies (Fig. 5). Furthermore, rhythmicity-based spectral segmentation reveals twice as many bands as previously described with two divergent modes of operation: activity in transient bands indicates neuronal inputs, whereas activity in high rhythmicity bands signifies maintenance of neuronal activity via sustained oscillations. This duality can conceivably expand the capacity and efficacy of neuronal computations and, with a spectrum spanning several orders of magnitude ^5^, can equip the brain with the flexibility necessary for a wide range of cognitive functions.

**Figure 9.**
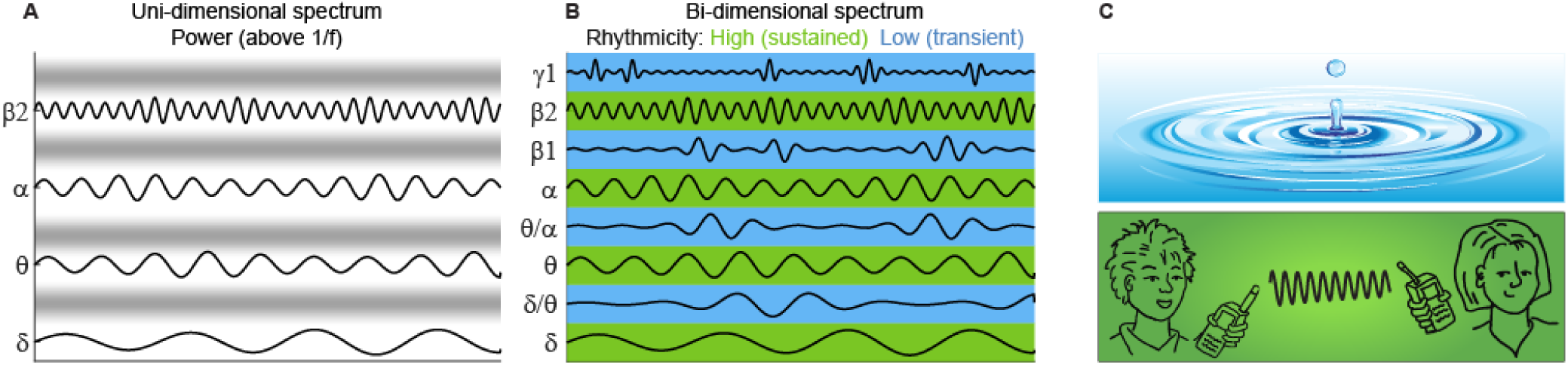
Rhythmicity reveals a bi-dimensional architecture. Brain rhythms can be classified along a new dimension of rhythmicity, revealing a universal spectral architecture: high-rhythmicity (sustained) bands maintain ongoing activity, while low-rhythmicity (bursty) bands denote the response to change. This framework generalises across species, sexes, recording techniques, and cognitive states. **A.** Traditional view – the spectrum is divided to logarithmically progressing oscillatory bands. Role of activity expressed in the gaps between band (blurred grey borders) is unclear. **B.** Proposed view – rhythmicity dimension distinguishes sustained (green) vs. bursty (blue) bands. **C.** Functional division between the two rhythmicity modes. Bursty bands are associated with the processing of transient incoming inputs (metaphorized as ripples resulting from a drop of water). Sustained bands enable the maintenance of brain states and the putative orchestration of engaged brain areas (metaphorized as a telecommunication through sustained radio frequency, green). Artwork credit: droplet adapted from Allvectors.com under CC BY 4.0 licence; telecommunication contributed to the manuscript by Mirjam Karvat.

Segmenting the spectrum according to rhythmicity also opens up theoretical avenues for understanding how different rhythmic regimes may support distinct neural coding strategies. For example, the bi-dimensional organization of spectral phenomena can bridge two prominent views of the brain’s *modus operandi*, namely *phase-coding* ^49^ and *rate-coding* ^50^. These views differ regarding whether the precise timing of neuronal outputs encodes information about neuronal inputs. *Phase-coding* posits that oscillations temporally bias neuronal responses to inputs: synaptic inputs that arrive during the excitable phase of an oscillation are more likely to cause outputs; whereas inputs arriving at the opposite phase get dismissed or curtailed ^51,52^. According to this view, oscillations causally influence the formation, synchronization, and sequence of cell assemblies, communication efficacy, and ultimately, an organism’s behaviour ^49,53–55^. Within the rhythmic architecture framework, this suggests that sustained oscillations may maintain temporal scaffolds for coordinating neuronal responses. Much like radio signals rely on stable carrier frequencies to transmit information, sustained rhythms may facilitate local and distant communication by aligning excitability windows across neuronal populations and networks ^56,57^.

In contrast, *rate-coding* proposes that the rate at which neurons fire contains all relevant information. According to one interpretation of rate-coding ^50^, enhancing neuronal firing-rate suffices for assembly formation, deeming oscillations unnecessary. In the context of rhythmic architecture, previous studies have shown that surges in firing rate tend to align with troughs of field potentials during oscillatory bursts—initially observed in the gamma-^23,58^ and beta-bands ^59^ and more recently across the entire 3– 120 Hz spectrum ^13^. Thus, even if oscillations are not required for information transfer per se, they serve as a valuable indicator of moments of heightened neuronal activity. Much like ripples on a calm water surface mark the impact of a falling object, oscillatory bursts—confined to low-rhythmicity frequency bands—act as spectral markers of cognitively relevant events.

An important advantage of introducing rhythmicity as an additional dimension is that it allows us to build upon, rather than discard, prior knowledge. We do not suggest that all transient bands, or all sustained bands serve the same function. There remains considerable scope for functional specialisation among specific bands. For example, within sustained rhythms, alpha activity is often associated with the maintenance of a widespread, general rest state ^63^, whereas beta2 has been linked to keeping more targeted forms of cognitive control ^61^.

Turning to transient bands, these may reflect inputs from distinct sources. For instance, evoked responses—often considered the frequency-domain analogue of time-domain event-related potentials— are typically dominant just below 10 Hz ^64^, suggesting that the theta/alpha transient band may reflect broad, stimulus-driven synchronisation. In contrast, beta bursts have been associated with the spontaneous activation of neural networks, where coincident bursts across regions may indicate functional connectivity ^65^. In this context, beta1 may be conceptualised as reflecting internally generated inputs, such as packets of action potential sent from one brain region to its targets ^66^. These examples highlight the potential of rhythmicity-based segmentation to support a more nuanced functional characterisation of the electrophysiological spectrum in future studies.

Our consistent observation that broad frequency bands can be subdivided into rhythmically distinct sub-bands may help clarify longstanding ambiguities about their functional roles. For instance, activity within the canonical beta band (∼12–30 Hz) has been associated with both sustained and burst-like dynamics. When conceptualised as sustained oscillations, beta is thought to support the maintenance of ongoing motor or cognitive states—for example, continuous beta activity over sensorimotor cortex has been linked to tonic motor inhibition or postural maintenance ^60^, and sustained beta synchronisation in frontoparietal networks has been associated with stabilising cognitive sets and top-down control, suggesting a role in signalling the cognitive status quo ^61^. Conversely, when viewed as transient bursts, beta activity has been linked to event-related processes, such as brief motor cortex beta bursts marking movement termination ^16^ or successful inhibition in stop-signal tasks ^62^. Notably, one study ^11^ demonstrated that both transient and sustained beta responses can co-occur within individuals, varying with task stage—transient responses following informative cues, and sustained responses during anticipatory periods. These were also spectrally distinct, occupying different sub-bands within the broader beta range. It is therefore plausible that previously reported, and at times seemingly contradictory, functional roles attributed to beta activity may in fact reflect spectral segregation into low- and high-rhythmicity sub-bands within the band referred to as beta.

A further promising direction for future research concerns the interactions between neighbouring frequency bands. We observed that high-rhythmicity bands can suppress the emergence of oscillations in adjacent bands via phase interference (Fig. 3A). Specifically, ongoing oscillatory activity in a dominant high-rhythmicity band may curtail the development of nearby rhythms through phase-based interactions (i.e., interference). This interpretation is supported by our finding that removing the high-rhythmicity band using a stop-band filter abolished the observed phase shift (Fig. 3A), consistent with LAVI’s sensitivity to phase differences. This interpretation, together with the dominance of high-rhythmicity events (higher power even when normalized to compensate the 1/f slope, Fig. 3F) also carries intriguing implications for the broader operational principles of the brain. It raises the possibility that high-rhythmicity bands may serve to establish dedicated “quiet” spectral zones—frequency ranges held in readiness for incoming input. In such a scenario, an external or internal input would produce a brief, ephemeral response, which is then rapidly curtailed by the surrounding rhythmic context. This mechanism would not only support a top-down role for sustained oscillations but also suggest that such control is not purely inhibitory. Rather, it may enhance contrast and selectivity by dynamically shaping the spectral landscape to prioritise response-relevant signals.

While our findings offer compelling and consistent insights into the functional architecture of brain rhythms, it is important to acknowledge certain limitations that may inform future refinements and interpretations. First, for practical reasons, we have chosen to focus on the 3–45 Hz range. This choice limited our ability to characterise dynamics in both lower (notably δ and δ/θ) and higher (γ and higher frequencies) frequency bands (Fig. 4). Nonetheless, both LAVI and ABBA are applicable to these ranges, provided that specific considerations are taken into account. For low frequencies, LAVI’s noise levels are influenced by session duration (fig. S6); thus, for frequencies below 3 Hz, we recommend using recordings of at least several minutes. Conversely, high-frequency LAVI is sensitive to the sampling rate, and we advise a minimum sampling frequency of 1 kHz. Additionally, power line noise (50/60 Hz) is highly rhythmic and can substantially affect frequencies above ∼40 Hz. Strong notch filters are not recommended, as they distort the rhythmicity of neighbouring frequencies (Fig. 3I, blue traces). Instead, we suggest using approaches such as spectral interpolation ^67^, which minimise distortion in the time domain while effectively reducing line noise.

Second, while our TMS findings provide causal support for the proposed spectral architecture, we acknowledge that neurostimulation studies inherently involve greater variability and interpretative complexity than observational analyses. Although the current protocol yielded informative results consistent with previous reports of spectrally confined responses to single-pulse TMS ^48^, it was constrained by factors such as sample size and the absence of behavioural correlates. Future studies could strengthen these findings by incorporating larger and more diverse cohorts, as well as concurrent behavioural or task-based measures to directly link rhythmicity modulation with cognitive function. Such refinements would enable a more precise characterisation of how rhythmic and arrhythmic inputs interact with distinct spectral modes and their functional relevance.

Despite these methodological limitations the present study points toward exciting future directions for investigating the basis and function of neural rhythmicity. The origins of neural oscillations—though extensively explored in both experimental and computational domains—can now be explored through the prism of distinct rhythmicity modes. A promising direction for future research lies therefore in exploring how the rhythmic architecture described here might reshape our understanding of the anatomical and computational bases of oscillatory activity. For instance, could certain anatomical structures be more conducive to one rhythmicity mode over another ^19^? Might distinct mechanisms underlie the generation of oscillations in the two modes—such as PING (pyramidal-interneuron gamma) circuits for bursty activity ^68^, versus pacemaker cells for sustained rhythms ^69^? (For a comprehensive review, see ^70^.) Furthermore, identifying the computational principles that best account for these two modes of operation—and understanding their functional and cognitive consequences—represents an exciting avenue for future investigation.

In conclusion, our findings introduce rhythmicity as a fundamental dimension of brain dynamics, supporting a dual sustained–bursty spectral architecture that underpins flexible and efficient neural computation. This framework generalises across species, sexes, recording modalities, and cognitive states. The practical implications of these theoretical insights are considerable: an individualised, detailed, and statistically robust spectral segmentation can be harnessed to inform brain stimulation protocols, enable non-invasive diagnostics for neuropathology, and guide the development of novel brain–computer interfaces.

## Methods

### Participants and data acquisition

We sought to establish the rhythmicity architecture as a universal phenomenon, therefore we analysed datasets from different laboratories and recording techniques. Details of the original publications from which data were analysed in this study is presented in the main text, Table 1.

### EEG acquisition and preprocessing

We performed all offline preprocessing and analysis steps using the Fieldtrip toolbox, release 20230822 ^71^ and custom code in Matlab 2020b (MathWorks, Natick, MA). In datasets I-III ^22,24^, data were sampled at 512 Hz using a g.GAMMAcap (gTec, Schiedlberg, Austria) and a g.HIamp amplifier (gTec). The cap had 62 electrodes distributed over the scalp, with the addition of two active earlobe electrodes. All electrodes were re-referenced offline to the average of the earlobe electrodes. In dataset IV ^25^, data were sampled at 2048 Hz using an ActiveTwo EEG system (Biosemi, Amsterdam, Netherlands). The cap had 32 electrodes that were re-referenced offline to the average of two electrodes over the temporal lobe (T7 and T8)^1^. Data in dataset V ^26^ was recorded with a Neuroscan (Compumedics, El Paso, TX, USA) NuAmps amplifier, with 30 channels sampled at 1000 Hz and re-referenced to electrodes over he left and the right mastoid. In dataset VI ^27^, data were acquired from 128 channels using EGI Hydrocel Geodesic Sensor Net (HGSN-128, Magstim EGI, Eugene, OR), sampled at 1000 Hz, and re-referenced offline to the average of all channels.

All succeeding preprocessing procedures were identical for all EEG datasets. We detrended, demeaned and bandpass filtered the data at 0.5-130 Hz. Then, we inspected the data visually and rejected bad channels, inserting interpolation of neighbouring channels instead of the bad channels. On average, we rejected 8.54% of electrodes from dataset I (SD 1.65%), 0.04% (0.26%) from dataset II, 2.3% (3.2%) from dataset III, 1.17% (2.22%) from dataset IV, 4.5% (1.9%) from dataset V, and 5.9% (5.0%) from dataset VI. Importantly, channel Cz, which was used for rhythmicity and oscillatory bursts analyses, was good for all 178 EEG subjects. Scalp muscle artefacts were detected and marked as epochs in which the amplitude of the signal at 100-120 Hz at all channels exceeded a threshold of 30 standard deviations. Finally, independent spatio-temporal components containing eye movements (blinks and saccades) were removed using independent component analysis (ICA). We preformed all preprocessing steps, as well as rhythmicity (LAVI), power, and oscillatory bursts analyses, on the whole session (that is, without dividing into trials).

For ECoG (dataset IX) we used all 907 channels reported “clean” in the original publication ^29^. In the per-area analysis (Fig. 4D-F) we excluded electrodes reported as “Medial” due to low count (n=12), or those that were marked as depth electrodes, and therefore lacked a cortical area assignment (n=28). After exclusion, this analysis consisted of 867 electrodes. For SEEG (dataset X ^30^) we used 733 electrodes reported as located in mesial-frontal, lateral-frontal, insula, medial-parietal, lateral-parietal, lateral-temporal, or mesial-temporal areas. In addition, for each area, we included the nearest concurrently recorded EEG scalp electrode. For LFP recordings, we used one STN channel from each hemisphere On and Off PD medication in dataset XI ^31^ sampled at 2048 Hz using a TMSi Porti (TMS International, Netherlands) or from the hippocampus of each rat in dataset XII ^32^.

### MEG acquisition and preprocessing

MEG data came from Stage 2 of the Cambridge Centre Aging and Neuroscience (Cam-CAN; www.cam-can.org; ^72^) study of healthy adult ageing (aged 18-88 years). Ethical approval was obtained from the East of England-Cambridge Central Research Ethics Committee, and participants gave full informed consent. A detailed description of exclusion criteria can be found in ^72^, Table 1. Of these, only participants with resting state MEG data that could be corrected for motion using MaxFilter were used here (n=625).

MEG data were collected using a 306-channel VectorView MEG system (Elekta Neuromag, Helsinki), consisting of 102 magnetometers and 204 orthogonal planar gradiometers, located in a magnetically shielded room. MEG resting state data (sampled at 1 kHz with a high-pass filter of 0.03 Hz) were recorded approximately 8.5 mins, while participants remained still in a seated position with their eyes closed, but instructed to stay awake. Head position within the MEG helmet was estimated continuously using four Head-Position Indicator (HPI) coils to allow offline correction of head motion.

The MaxFilter 2.2.12 software (Elekta Neuromag Oy, Helsinki, Finland) was used to apply temporal signal space separation ^73^ to the continuous MEG data to remove noise from external sources (correlation threshold 0.98, 10-sec sliding window), to continuously correct for head-motion (in 200-ms time windows), to remove mains-frequency noise (50-Hz notch filter), and to detect and reconstruct noisy channels. Following these de-noising steps, data were imported into Matlab using SPM12 (http://www.fil.ion.ucl.ac.uk/spm). These data are available on request from https://cam-can.mrc-cbu.cam.ac.uk/dataset/.

The first 20 seconds of data were ignored to allow participants to settle, and any samples after 542 seconds were ignored, in order to match data length across participants (to the minimum duration across participants). These data were then extracted from 4 magnetometers around Cz.

### Lagged Angle Vector Index (LAVI)

We measured rhythmicity as the consistency of the relations of phase (angle) between data points separated by a fixed number of cycles (“lag”, with the corresponding duration in ms determined by each frequency, and see Fig. 1 for considerations for choosing lag duration). The rational for defining rhythmicity in this manner is that the phase of a sustained oscillation at any time-point should predict its future phase ^18^.

We computed the time-frequency representation *x̂* for each time-point *t* and frequency of interest *f* using complex Morlet wavelets, of the form

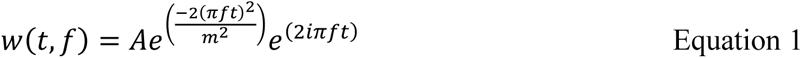

where *i* is the imaginary unit and *m* is the wavelet width, measured in cycles. The normalization factor was 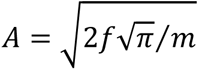. The value of *m* was set to 5, based on the exploration presented in Fig. 1E and fig. S2D. Care was taken to produce angles with the following convention: cosine must always be 1 and sine must always be cantered in upgoing flank, so the centre of the wavelet has the angle of 0 rad ^71^. The time-frequency representation was computed by convolving the signal with the Morlet wavelets at each frequency. For computational efficiency, this was implemented in the frequency domain by multiplying the Fourier transform of the signal with the Fourier-transformed wavelets, and applying an inverse Fourier transform to obtain the time-varying complex spectra at each frequency. Then, the rhythmicity λ was measured for each frequency as the magnitude of the vector mean vector of all angle differences (“coherence”) between the original time-frequency representation 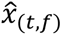 and a copy of itself with a constant lag of τ cycles 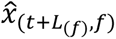:

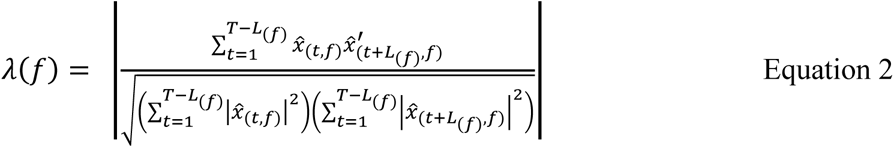

where *T* is total number of time points in the session, *L_(f)_* is the number of time points in τ cycles and the superscript ′ denotes the complex conjugate transpose. By definition, *λ* assumes values between 0 and 1. The median of *λ* (over *f*) is stable across subjects and depends on *m* and τ. Based on the exploration presented in Fig. 1D and fig. S2D, we set τ to 1.5 cycles, which together with *m* = 5 cycles yields a median λ of ∼0.4, providing a good dynamic range (for further discussion, see Supplementary Text, Parametrical effects on rhythmicity measures).

### Automated Band Border Algorithm (ABBA)

Using the LAVI median as a threshold, one can define areas above the median as “sustained” bands, and areas below as “transient” bands. However, fluctuations above and below the median are present also in the rhythmicity profiles of purely random signals (noise). The median of the rhythmicity profiles of noise also depends of *m* and τ, and their dispersion range depends on the sampling frequency, session duration, and the aperiodic slope (for further discussion, see Supplementary Text, parametrical effects on rhythmicity, and fig. S6). This relationship can be leveraged to determine spectral bands with statistical confidence using a bootstrap approach: frequencies whose rhythmicity levels exceed (or fall below) the top or bottom k observations from n bootstrap samples of noise can be considered sustained (or transient), corresponding to a two-tailed Type I error rate of α = k/n. In our analysis, we generated n = 200 surrogate noise samples and identified frequencies as sustained or transient if their rhythmicity levels fell within the top or bottom 2.5% (k = 5) of the bootstrap distribution (α = 0.05).

To generate the distribution of rhythmicity profiles we generated surrogates that matched the aperiodic component (1/*f*) of each subject. We calculated the aperiodic component of the EEG by fitting an exponential function to the EEG power spectrum, and used the fitted values as the input for an Iterative Amplitude Adjusted Fourier Transform algorithm (IAAFT, ^74–76^). IAAFT generates surrogate time series with a desired power spectrum (in the frequency domain) and data values (in the time domain) by keeping the original magnitude of the Fourier coefficients and assigning random phases. In each iteration of the algorithm there are five steps: a. initializing with a randomly shuffled version of the original time series to destroy temporal correlations while preserving the amplitude distribution; b. computing the Fourier transform of the shuffled time-series; c. replacing the magnitude of the Fourier coefficients with the desired magnitudes (i.e., those of the original time series) while keeping the current (random) phase; d. computing the inverse Fourier transform to obtain a new time series with the desired spectral properties, and e. matching the amplitude distribution: this is done pointwise by ranking the new time series and replacing its values with those of the original series sorted by rank (e.g., the highest value in the new series receives the highest value of the original series, the second-highest receives the second-highest, etc.). Since step (e) alters the spectral properties, this algorithm is repeated iteratively until the error between the desired and current spectra falls below a defined threshold (here, 2×10⁻⁴ of the original standard deviation).

The IAAFT implementation for Matlab ^77^ can take the original EEG values as input, but since noise levels can be predicted from experimental parameters (Supplementary Text—Parametrical Effects on Rhythmicity, and fig. S6) and in order to save computational time and facilitate the computation of hundreds of channels, we generated surrogate distributions of 200 repetitions with different sampling frequencies, durations, and aperiodic exponents. This resulted in a look-up table that was used to define significance limits for each frequency and channel with α = 0.05. Then, ABBA defines each frequency with a LAVI value above (below) the significance limit as significantly sustained (transient). Frequencies between points of crossing the median are then grouped into bands. The local maximum is taken as the peak frequency of sustained bands, and the local minimum is the peak frequency of transient bands. Since most subjects have a clear peak in the rhythmicity profile at the alpha band, the peak between 6 to 14 Hz can be used as an anchor to automatically allocate bands identity: the frequency with highest rhythmicity between 6 and 14 Hz is taken as alpha, the trough following alpha, as the frequency increases (i.e., to the right of alpha) is beta1, the next peak is beta2, and so forth, and also for frequencies lower than alpha. A Matlab implementation for LAVI and ABBA (including pink surrogate and look-up table generation) is available at https://github.com/laaanchic/LAVI. This code is using functions provided by Fieldtrip ^71^ and Venema V. ^77^, and is shared under the GNU General Public License (GPL) and Berkeley Software Distribution (BSD) license, respectively.

In fig. S2 we compare band-detection with LAVI and ABBA to a commonly used power-based method. To this end, we computed the power spectrum of the EEG of 90 participants (using the Matlab function ‘pwelch’), and used Spectral parameterization (specparam, formerly fooof ^8^) with recommended default configurations to detect the range and peak frequency of bands in the frequency range 2-40 Hz.

### Simulations

For all simulations presented in Figs. 3, S1, S4, and S5, we generated 180 s “pink” surrogate data using IAAFT, with aperiodic exponent of −1, range of ∼100 μV, mean 0 μV and SD ∼12.5 μV. To manipulate the activity in a specific frequency band, we first decomposed the original signal using an array of third-order Butterworth bandpass filters (centred at 3–100 Hz in 1-Hz steps, each with a 1-Hz bandwidth). This decomposition was performed once and served as a common basis for all subsequent manipulations. In each manipulation iteration, we selectively modified the signal in the 15-Hz band and then reconstructed the full signal by summing all band-limited components. Finally, we computed and stored the rhythmicity profile (LAVI) of the reconstructed signal using the same parameters applied throughout the manuscript (wavelet width of 5 cycles and a lag of 1.5 cycles).

To simulate “bursts” of varying durations in an otherwise quiet band (Fig. 3H-I), we first attenuated the signal filtered at 14–16 Hz by multiplying it by 0.5. We then introduced bursts by multiplying segments of the signal by 2, with a smooth rise and fall. Burst duration was the independent variable, ranging logarithmically from 2 to 20 cycles, while the inter-burst interval was jittered between 5 and 15 cycles.

To simulate bursts with varying degrees of frequency drift (Fig. S4A–B), we defined each burst as a sinusoid of the form y = sin(2π *f t*), where the instantaneous frequency *f* linearly drifted from 15+drift/2 to 15-drift/2 Hz. The slope of this drift was the dependent variable and ranged logarithmically from 0.1 to 10 Hz.

In the transient input simulations (fig. S4C-F), the initial conditions were identical to the oscillatory burst duration simulation. Each transient input was simulated as an impulse (one sample long) with amplitude equal to the EEG range (100 μV). The number of impulses in a “burst” was the independent variable and ranged linearly from 1 to 13. In the rhythmic condition (fig. S4C-D), the distance between each impulse in a “burst” was set to 66.6 ms (one cycle of a 15 Hz oscillation). In the arrhythmic condition (fig. S4E-F), the last impulse in a “burst” was set to 66.6*(*n*-1) ms after the first pulse, with *n* the number of impulses in a “burst”. Then, the remaining *n*-2 impulses were randomly distributed between the first impulse and last impulse. Like in the burst duration simulation, the inter “burst” interval jittered between 5 and 15 cycles.

To implement phase shifts (fig. S1), we flipped the sign of the signal by multiplying it by −1 every 2 to 20 cycles (in logarithmic steps). To investigate the effect of power on rhythmicity (fig. S5A-B), we multiplied the data filtered at 15 Hz by values ranging between 0.1 and 2, in steps of 0.1. In all simulations, we defined the significance limits as the maximum and minimum rhythmicity values of the original surrogate data, and the main dependent variables were the rhythmicity values at the manipulated frequency (15 Hz). All rhythmicity calculations were made with Wavelet width of 5 cycles and lag of 1.5 cycles.

As can be observed in Fig. 3I, fig. S1B, and fig. S5B, increasing (decreasing) the rhythmicity in one band decreases (increases) rhythmicity in neighbouring bands. To investigate if the troughs in the rhythmicity profiles we observed in the data of 90 subjects can be explained as an (artefactual) effect of the peak in alpha, we generated 90 instantiations of 120 s 1/*f* surrogates. We decomposed the surrogates into discrete frequencies, and multiplied the amplitude of the signal in 11 Hz (alpha) by a factor randomly chosen between 1.05 and 2.25. We then calculated the rhythmicity profiles (LAVI) and detected bands (ABBA). Then, we calculated the fit between the LAVI values at alpha peak and beta1 trough of the surrogate population with a linear regression. We used the coefficients of the regression to calculate the expected beta1 troughs based on data alpha peaks, and compared them to beta1 troughs observed in data (fig. S5C-D). Finally, we compared the LAVI values at theta/ alpha and beta1 troughs of the data to the troughs of surrogates generated with aperiodic components and power in alpha matching the observed data, using filter arrays (fig. S5E).

### Bursts analysis

To detect oscillatory bursts, we first estimated the power of the raw EEG using Morlet wavelets with a width of 5 cycles, centred at frequencies spanning logarithmically from 3 to 45 Hz. Artefactual segments—defined as time points where the EEG exceeded 250 μV or intervals marked as muscle artefacts during preprocessing—were removed, along with a 500 ms buffer before and after each segment ^78^. Burst peak time and frequency were identified as local maxima in the two-dimensional time–frequency plane that exceeded the 90^th^ percentile of power. We then recorded the peak frequency and its corresponding time point.

To align individual bursts to a common reference point and compute the burst-related potential (BRP; Fig. 3A), we band-pass filtered the raw signal using the band borders defined by the LAVI and ABBA methods as the filter’s corner frequencies. We then identified the nearest trough in the filtered signal and defined it as time zero (t = 0), extracting an epoch of ±3 cycles around this point. To assess potential inter-band influences, we also extracted data at the time of each burst while suppressing activity in neighbouring bands (with notch filters).

Next, we determined the burst onset and offset in two steps. First, we defined the initial burst boundaries as the time points at which power at the peak frequency dropped below the 75^th^ percentile. Next, we identified all frequencies that peaked at any time point within this initial window. The final burst onset and offset were then defined as the time points at which all of these peak frequencies fell below the 75^th^ percentile of either power (Fig. 3B) or rhythmicity (WTPL, Fig. 3C).

Since we were interested in average durations and rates of bursts in the different bands over the whole session, we paid special attention to avoiding bursts overlapping in time/frequency. Therefore, if two bursts overlapped in time, and also had a peak frequencies difference smaller than a quarter of the peak frequency of either burst, they were merged into one burst. The merged burst was assigned the frequency of the burst with the higher energy (estimated as power x duration).

Then, to calculate the burst rate (Fig. 3D), we counted the total number of bursts in each band across the entire session and divided by the session duration and band width. This latter normalisation accounts for the fact that the band width increases logarithmically with frequency.

Occupancy (Fig. 3E) was calculated by summing up all time points between the beginning and end of each burst in each band, dividing by the total number of samples, and multiplying by 100 to obtain percentages.

Relative power (Fig. 3F) was defined as the dB value (10 × log_10_) of the ratio between the peak-power and the 90^th^ power percentile of the peak-frequency.

Finally, band consistency (Fig. 3G) was defined, for each burst, as the number of time-points in which the peak-frequency was confined within the LAVI-defined frequency band of the burst itself, divided by the total number of samples in the burst, and multiplied by 100 for percentage representation.

### TMS-procedure

All procedures were approved by the Cambridge Psychology Research Ethics Committee. All participants provided written informed consent prior to data acquisition for the study. Biphasic single and repetitive TMS pulses were delivered using a DuoMAG XT-100 TMS stimulator and a figure-of-eight coil DuoMAG 70BF (Brainbox Ltd, Cardiff, UK). During the stimulation, participants sat in a comfortable recliner chair with a neck rest. To ensure the precise targeting of specific brain regions, the coil was controlled via Brainsight 2 neuronavigation system (Brainbox) in combination with an Axilum TMS-Cobot (Axilum Robotics, Schiltigheim, France). The Cobot is a robotic system that actively monitors and adjusts the positioning of the coil, and compensates for head movements throughout the experiment. To detect head movements, participants wore a headband reference tracker that was monitored by a Polaris Vega ST camera (NDI, Waterloo, Canada). The TMS coil was oriented with the handle pointing posteriorly with respect to the participant’s head, at an angle of 45 degrees relative to midline. The MNI coordinates for dorso-lateral prefrontal cortex (dlPFC, x = 33, y = 39, z = 26) stimulation were derived from the peak voxel showing the strongest effect in BOLD signal in a previous meta-analysis study ^79^.

Our protocol established a TMS stimulation intensity at 90% of the resting motor threshold (RMT) of each participant. To determine the RMT, we positioned the coil over the hand area of the right primary motor cortex and asked the participant to keep their left hand relaxed and at rest. Then, we determined the minimum intensity at which a single TMS pulse produced a visible twitch in the abductor pollicis brevis muscle of their left hand, in five of ten successive pulses.

EEG recordings were obtained with the actiCHamp Plus 64 system (Brain Products GmbH, Gilching, Germany), which is TMS-compatible. The system includes a DC-coupled amplifier avoiding AC recoding and high-pass filter during the recording period. EEG signals were acquired from 64 active electrodes arranged on an actiCAP slim electrode cap. The ground electrode was placed at FPz, and the reference at Cz. Electrode impedance was maintained below 20 kOhm. We used a sampling rate of 1000 Hz for the EEG resting-state recording and 5 kHz for the TMS-EEG recordings.

After EEG setup, we used LAVI and ABBA to determine the individual alpha, beta1, and beta2 peak frequencies that would define the repetitive TMS stimulation frequencies. For this, we performed a resting-state EEG recording of 12 minutes (2-minutes with open eyes, 8-minutes closed eyes, and 2-minutes open eyes). During the fixation periods, participants were instructed to keep their eyes still and look at a white fixation cross presented on a black background. LAVI was calculated over the closed-eyes period.

The TMS-EEG session comprised a total of 480 trials distributed into 8 blocks. There were 7 conditions: 1 single pulse (sp-TMS), 3 rhythmic conditions consisting of a 6-pulses train at alpha, beta1 or beta2 individual frequencies, and 3 arrhythmic conditions, also consisting of 6-pulses train. For each arrhythmic condition we precomputed patterns of 6-pulses that excluded frequencies within a +/-2-Hz band centred on the frequency of the corresponding rhythmic condition, their harmonics, subharmonics and over 50 Hz. The length of both rhythmic stimuli was equal to 5 cycles of the peak frequency. Each block included 12 trials of the single pulse condition and 8 trials of each of the rhythmic and arrhythmic conditions, presented sequentially in a pseudorandomized order. During each block, participants were instructed to keep their eyes still and look at a white fixation cross presented on a black background. The inter-train interval (ITI) between two consecutive trials was adjusted to the preceding frequency according to safety guidelines ^80^. For frequencies up to 10 Hz, we used a 3-s ITI; for frequencies over 10 Hz and up to 15 Hz, we used a 5-s ITI; for frequencies over 15 Hz and up to 20 Hz, we used an 8-s ITI; for frequencies over 20 Hz and up to 25 Hz, we used a 10-s ITI. Stimulus and TMS pulses delivery was controlled using Psychtoolbox-3 ^81^ in Matlab.

### TMS-data analysis

EEG data were pre-processed offline in Matlab using Fieldtrip and custom-written code, following previously published guidelines ^82,83^. First, we removed the TMS ringing artefacts by replacing data values from 1 ms before to 15 ms after the TMS pulse trigger with NaNs (Not-a-Number). Then, we down-sampled the data to 1000 Hz. Afterwards, we visually inspected the data and marked fragments with major artefacts (e.g., jumps, movement, muscle); these fragments were also replaced by NaNs. Then, all the fragments with NaNs were interpolated with a linear method. To detect remaining TMS-related artefacts, we ran a first independent component analysis (ICA) with the FastICA algorithm. We averaged the components’ signal 50 ms after TMS pulse over all trials and rejected those components whose amplitude exceeded 30 μV. Then, we ran a second ICA. In this run, components relating to eye blinks, eye movements, muscle and sensor-localized noise were identified and removed. All bad data fragments were replaced via Piecewise Cubic Hermite Interpolation (pchip). Then, the Cz reference channel was recovered and the data was re-referenced to a common average reference. Finally, the data were cut into segments of each trial type, including a pre-train onset interval of 2.5 sec and post-train offset interval of 2 sec.

### Time-resolved rhythmicity

To measure rhythmicity-based burst-durations (Fig. 3C) and how rhythmicity changes in time within a trial in response to a visual stimulus (Fig. 7B) or TMS pulses (Fig. 8D and 8F), we used the Within Trial Phase Lock (WTPL) method. This method is described in detail elsewhere ^22^. In brief, WTPL measures time-resolved rhythmicity by asking how sustained the oscillation is for at least one period. Like LAVI, WTPL assumes that sustained oscillations are repetitive, and as such, the phase at any time point is expected to predict future and past phases. Specifically, the phase one cycle before or after a time point should be near zero, whereas phase slips cause deviations of this phase relation away from zero. In each frequency of interest, time-point, and trial, we computed the difference between the phase in the time-point of interest (*ϕ* _0_), one cycle beforehand (*ϕ* _-1_) and one cycle after (*ϕ* _1_).

The WTPL is computed as the mean resultant vector length of the two phase relations:

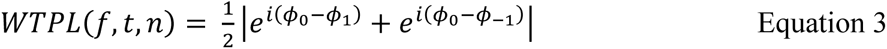

where *f* stands for frequency, *t* for timepoint, and *n* for trial. A Matlab implementation for WTPL is available at https://github.com/laaanchic/WTPL.

Note that since WTPL baseline levels of certain frequencies are higher than others (as learned from LAVI), between-frequency analysis is done after baseline subtraction (termed ΔWTPL in Fig. 7 and Fig. 8). We took the duration −1 s to −0.5 s (relative to the first TMS impulse) as the baseline. Note, that the TMS frequency used for each participant was dependent on their individual band peak frequency, which varied across participants. Therefore, in Fig. 8, we plotted the population-averaged ΔWTPL values after normalizing individual frequencies (that is, setting the peak frequency per band to “0”).

### Amplitude covariance

We used the High-Frequency (HF) activity estimation and separation of electrodes into positively and negatively responding to the visual stimuli as reported in ^29^. To estimate HF, the entire signal was band-pass filtered into eight 10 Hz sub-ranges between 70 and 150 Hz. Then, instantaneous amplitude in each band was extracted using the Hilbert transform. To account for the 1/*f* profile of the power spectrum, the amplitude in each sub-range was normalized by dividing the instantaneous amplitude by the mean amplitude in that range. Finally, the amplitude traces from all sub-bands was averaged. Channels were defined as responsive if the HF activity in one of four non-overlapping “stimulus-on” windows (0.1-0.3 s, 0.3-0.5 s, 0.5-0.7 s, and 0.7-0.9 s relative to stimulus onset) differed significantly from baseline activity (−0.2-0 s) for at least one of the four categories of visual stimuli used in the study (faces, watches, objects, and animals). Trials containing excessive noise between −0.3 and 1.6 s relative to stimulus onset, or with stimulus duration shorter than 900 ms, were excluded from analysis. For each category, the mean HF signal during the “stimulus-on” window was compared to the mean HF during baseline (two-tailed paired *t*-test with Benjamini-Hochberg false discovery rate correction ^84^ across electrodes and Bonferroni correction across windows). Hence, electrodes with qFDR < 0.05/4 in at least one of the four windows were considered responsive. Electrodes were considered “positively responsive” if activity during the “stimulus-on” window increased relative to baseline, and “negatively responsive” when it decreased.

To estimate the Low-Frequency (LF) activity, we computed the time-frequency representation of the raw data using complex Morlet wavelets. The wavelets were centred on frequencies spanning logarithmically from 3 to 45 Hz, with width of 5 cycles each. To extract the amplitude, we took the complex magnitude (absolute value) of the complex wavelet output. This method was chosen to maintain consistency with the LF analysis used throughout the manuscript, including for LAVI computation. In contrast, the HF estimation followed the approach used in the original publication ^29^. We then computed for each electrode and each Low-Frequency the covariance between the LF amplitude and normalised (*z*-transformed across time) HF and activities. This normalization was performed to account for differences in HF activity between electrodes. We calculated the covariance over the whole session (grey trace in Fig. 7C) or in response to the visual stimulus onset (0-0.2s, purple traces in Fig. 7C).

We computed the time-frequency representation *x*^ for each time-point *t* and frequency of interest *f* using complex Morlet wavelets (Equation 1).

### Statistical tests

All statistical tests were performed using Matlab with the Statistics and Machine Learning toolboxes. Data throughout this manuscript are presented as mean ± *SEM* unless otherwise stated. Significance level was set at α = 0.05. If multiple tests were performed (e.g., different measures in Fig. 3), their *p*-values were corrected using the false discovery rate according to the Benjamini-Hochberg algorithm ^84^. For post-hoc tests following ANOVA, we used the Fisher’s least significant difference procedure. All statistical tests are two-sided. Complete details of all statistical tests are presented in Table S1.

To define bands with statistically significant increased or decreased rhythmicity we adopted a bootstrap approach using a look-up table with α = 0.05 (see above, [Automated Band Border Algorithm (ABBA)]).

To test for differences in band-peak distributions between datasets or areas (Fig. 4), as well as for differences in burst characteristics between bands (Fig. 3), we used a one-way ANOVA (Matlab function “anova1”). In figures 3 we excluded participants with > 3 SD, resulting in the exclusion of 1 participant.

To assess whether two or more distributions differed, we computed the likelihood of the data under both the null and alternative hypotheses using Bayes Factors, which quantify the relative evidence for or against a difference. We used Bayes Factor (BF) t-tests or ANOVA, using a BF toolbox for Matlab ^85^. The output of the bf.ttest or bf.anova Matlab function is BF_10_, or how strong is the evidence to support H_1_ (the distributions are different) and reject H_0_ (the distribution are not different). Its counterpart BF_01_=1/BF_10_ estimated how strong is the evidence to support H_0_ and reject H_1_. To determine the strength of this evidence we followed the guidelines suggested in ^86^: 3<BF_10_<10 indicates a moderate evidence for H_1_, and BF_10_>10 indicates a strong evidence for H_1_. Similarly, 1/10<BF_10_<1/3 (or 3<BF_01_<10) indicates a moderate evidence for H_0_, and BF_10_<1/10 (or BF_01_>10) indicates a strong evidence for H_0_. Default Cauchy priors for effect sizes were used (scale parameter = 0.707). No Markov-chain Monte-Carlo sampling was required, as Bayes factors were calculated analytically.

For comparing LAVI values across frequencies (Figs. 2D and 6C) we adopted a cluster-permutation approach ^87,88^. This non-parametric method is appropriate to control the family-wise error rate since values of neighbouring frequencies are not independent from each other. We normalized the values of each subject by subtracting the median across frequencies. Then, we computed the *t* test at each frequency. After computing *t* values, we clustered neighbouring points exceeding a significance level of α = .05 and summed the *t* values of each cluster. This sum served as the value for comparison in the cluster-level statistics. We then created 1000 permutations of data with shuffled labels (i.e., eyes open or closed in Fig. 2D or medication On or Off in Fig. 6C), and took the *t* sum of the largest cluster in each permutation. We considered clusters in the original data as significant if their summed *t* values were higher than 99.5% or lower than 0.5% of the shuffled distribution (that is, *p*<0.01, two-tailed) or higher than 97.5% or lower than 2.5% (*p*<0.05), as specified in the main text. Cluster-permutation analysis of amplitude covariation (Fig. 7C) was calculated using similar parameters, but here the *t*-test was performed between trial data (purple traces) and the covariance over the whole session (grey traces). Significant clusters of WTPL in time and frequency (Figs. 7B, 8D, and 8F) were defined with *t*-sums computed relative to a baseline before the visual stimulus or TMS pulses (−1 to −0.5 s), over 1000 repetitions (ECoG) or 64 repetitions (2^6^, maximal permutation possible with six TMS subjects).

## Data availability

The raw data of each dataset used in this study are available at:

I: https://figshare.com/articles/dataset/EEG_during_the_two-flash_task/29519285

II: https://figshare.com/articles/dataset/EEG_at_wakeful_rest/29519639/1

III: https://figshare.com/articles/dataset/EEG_during_tactile_timing_task/29519756/1

IV: https://dataverse.harvard.edu/dataset.xhtml?persistentId=doi:10.7910/DVN/YD7PPU

V: requests for data should be directed to and will be fulfilled by the corresponding authors of ref. *25*, Mikael Johansson and Robin Hellerstedt.

VI: further information and requests should be directed to and will be fulfilled by the lead contact of ref. *26*, Pierre Gagnepain.

VII: https://figshare.com/articles/dataset/EEG_at_rest_and_with_TMS/27934962/2

VIII: https://camcan-archive.mrc-cbu.cam.ac.uk/dataaccess/

IX: https://osf.io/4hxpw/

X: http://epi.fizica.unibuc.ro/scalesoldnew/.

XI: https://data.mrc.ox.ac.uk/stn-lfp-on-off-and-dbs

XII: https://portal.nersc.gov/project/crcns/download/hc-3

## Code availability

A Matlab implementation for LAVI and ABBA (including pink surrogate and look-up table generation) is available at https://github.com/laaanchic/LAVI.

## Footnotes

1 As recommended for channel level analysis when earlobe electrodes are unavailable, see https://www.fieldtriptoolbox.org/getting_started/biosemi/

## Acknowledgments

We thank Profs. Rik Henson and Leon Deouell for insightful comments as well as data use. We also thank Flor Kusnir, Nir Ofir, Anna Goodman, and Yarden Weiss, for providing helpful remarks on previous drafts of this article.

## Funding sources

German Research Foundation Walter-Benjamin post-doctoral fellowship KA 5804/1-1 (GK)

Azrieli Foundation graduate fellowship (GV)

Edmond and Lily Safra Center fellowship for Brain Sciences graduates (GV)

Medical Research Council grant MC-A060-5PR00 (MCA)

James McDonnell Scholar Award in Understanding Human Cognition (ANL)

ISF grant 958/16 (ANL)

European Research Council (ERC) under the European Union’s Horizon 2020 research and innovation programme 852387 (ANL)

FLAG-ERA collaborative grant (MONAD) and the Israel Innovation Authority.

## Author contributions

Conceptualization: GK

Data curation: GK, GV

Formal analysis: GK, MCG, GV

Funding acquisition: GK, MCA, ANL

Investigation: GK, MCG

Methodology: GK, ANL

Project administration: GK, MCA, ANL

Resources: MCA, ANL

Software: GK, MCG, GV

Supervision: MCA, ANL

Visualization: GK, ANL

Writing – original draft: GK, MCA, ANL

Writing – review & editing: GK, MCG, GV, MCA, ANL

## References

1. Buzsáki, G., Anastassiou, C. A. & Koch, C. The origin of extracellular fields and currents — EEG, ECoG, LFP and spikes. Nat Rev Neurosci 13, 407–420 (2012).

2. Buzsáki, G. Rhythms of the Brain. vol. xv (Oxford University Press, New York, NY, US, 2006).

3. Kane, N. et al. A revised glossary of terms most commonly used by clinical electroencephalographers and updated proposal for the report format of the EEG findings. Revision 2017. Clin Neurophysiol Pract 2, 170–185 (2017).

4. Başar, E. & Güntekin, B. A review of brain oscillations in cognitive disorders and the role of neurotransmitters. Brain Research 1235, 172–193 (2008).

5. Buzsáki, G. & Draguhn, A. Neuronal Oscillations in Cortical Networks. Science 304, 1926–1929 (2004).

6. Buzsáki, G. & Mizuseki, K. The log-dynamic brain: how skewed distributions affect network operations. Nat Rev Neurosci 15, 264–278 (2014).

7. Newson, J. J. & Thiagarajan, T. C. EEG Frequency Bands in Psychiatric Disorders: A Review of Resting State Studies. Frontiers in Human Neuroscience 12, (2019).

8. Donoghue, T. et al. Parameterizing neural power spectra into periodic and aperiodic components. Nat Neurosci 23, 1655–1665 (2020).

9. Han, C. et al. Multiple gamma rhythms carry distinct spatial frequency information in primary visual cortex. PLoS Biol 19, e3001466 (2021).

10. Nougaret, S. et al. Low and high beta rhythms have different motor cortical sources and distinct roles in movement control and spatiotemporal attention. PLOS Biology 22, e3002670 (2024).

11. Kilavik, B. E. et al. Context-Related Frequency Modulations of Macaque Motor Cortical LFP Beta Oscillations. Cereb. Cortex 22, 2148–2159 (2012).

12. van Ede, F., Quinn, A. J., Woolrich, M. W. & Nobre, A. C. Neural Oscillations: Sustained Rhythms or Transient Burst-Events? Trends in Neurosciences 41, 415–417 (2018).

13. Karvat, G., Alyahyay, M. & Diester, I. Spontaneous activity competes with externally evoked responses in sensory cortex. PNAS 118, (2021).

14. Sherman, M. A., et al. Neural mechanisms of transient neocortical beta rhythms: Converging evidence from humans, computational modeling, monkeys, and mice. PNAS 113, E4885–E4894 (2016).

15. Engel, A. K., Fries, P. & Singer, W. Dynamic predictions: Oscillations and synchrony in top–down processing. Nat Rev Neurosci 2, 704–716 (2001).

16. Feingold, J., Gibson, D. J., DePasquale, B. & Graybiel, A. M. Bursts of beta oscillation differentiate postperformance activity in the striatum and motor cortex of monkeys performing movement tasks. PNAS 112, 13687–13692 (2015).

17. Lundqvist, M. et al. Gamma and Beta Bursts Underlie Working Memory. Neuron 90, 152–164 (2016).

18. Fransen, A. M. M., van Ede, F. & Maris, E. Identifying neuronal oscillations using rhythmicity. NeuroImage 118, 256–267 (2015).

19. Myrov, V. et al. Rhythmicity of neuronal oscillations delineates their cortical and spectral architecture. Commun Biol 7, 1–18 (2024).

20. Rayson, H. et al. Detection and analysis of cortical beta bursts in developmental EEG data. Developmental Cognitive Neuroscience 54, 101069 (2022).

21. Berger, H. Über das Elektrenkephalogramm des Menschen. Archiv f. Psychiatrie 87, 527–570 (1929).

22. Karvat, G., Ofir, N. & Landau, A. N. Sensory Drive Modifies Brain Dynamics and the Temporal Integration Window. Journal of Cognitive Neuroscience 36, 614–631 (2024).

23. Atallah, B. V. & Scanziani, M. Instantaneous Modulation of Gamma Oscillation Frequency by Balancing Excitation with Inhibition. Neuron 62, 566–577 (2009).

24. Ofir, N. & Landau, A. N. Neural signatures of evidence accumulation in temporal decisions. Current Biology 32, 4093–4100.e6 (2022).

25. Noguchi, Y. Individual differences in beta frequency correlate with the audio–visual fusion illusion. Psychophysiology 59, e14041 (2022).

26. Hellerstedt, R., Johansson, M. & Anderson, M. C. Tracking the intrusion of unwanted memories into awareness with event-related potentials. Neuropsychologia 89, 510–523 (2016).

27. Legrand, N. et al. Attentional capture mediates the emergence and suppression of intrusive memories. iScience 25, 105516 (2022).

28. Taylor, J. R. et al. The Cambridge Centre for Ageing and Neuroscience (Cam-CAN) data repository: Structural and functional MRI, MEG, and cognitive data from a cross-sectional adult lifespan sample. NeuroImage 144, 262–269 (2017).

29. Vishne, G., Gerber, E. M., Knight, R. T. & Deouell, L. Y. Distinct ventral stream and prefrontal cortex representational dynamics during sustained conscious visual perception. Cell Reports 42, (2023).

30. Barborica, A. et al. Studying memory processes at different levels with simultaneous depth and surface EEG recordings. Front. Hum. Neurosci. 17, (2023).

31. Wiest, C. et al. Local field potential activity dynamics in response to deep brain stimulation of the subthalamic nucleus in Parkinson’s disease. Neurobiology of Disease 143, 105019 (2020).

32. Mizuseki, K., Sirota, A., Pastalkova, E. & Buzsáki, G. Theta oscillations provide temporal windows for local circuit computation in the entorhinal-hippocampal loop. Neuron 64, 267–280 (2009).

33. Ball, T., Kern, M., Mutschler, I., Aertsen, A. & Schulze-Bonhage, A. Signal quality of simultaneously recorded invasive and non-invasive EEG. NeuroImage 46, 708–716 (2009).

34. Nuñez, A. & Buño, W. The Theta Rhythm of the Hippocampus: From Neuronal and Circuit Mechanisms to Behavior. Front. Cell. Neurosci. 15, (2021).

35. Buzsáki, G. Theta rhythm of navigation: Link between path integration and landmark navigation, episodic and semantic memory. Hippocampus 15, 827–840 (2005).

36. Jacobs, J. Hippocampal theta oscillations are slower in humans than in rodents: implications for models of spatial navigation and memory. Philosophical Transactions of the Royal Society B: Biological Sciences 369, 20130304 (2014).

37. Buzsáki, G. & Moser, E. I. Memory, navigation and theta rhythm in the hippocampal-entorhinal system. Nat Neurosci 16, 130–138 (2013).

38. Vanderwolf, C. H. Hippocampal electrical activity and voluntary movement in the rat. Electroencephalogr Clin Neurophysiol 26, 407–418 (1969).

39. Grady, C. The cognitive neuroscience of ageing. Nat Rev Neurosci 13, 491–505 (2012).

40. Babiloni, C. et al. Sources of cortical rhythms in adults during physiological aging: A multicentric EEG study. Human Brain Mapping 27, 162–172 (2006).

41. Merkin, A. et al. Do age-related differences in aperiodic neural activity explain differences in resting EEG alpha? Neurobiology of Aging 121, 78–87 (2023).

42. Jenkinson, N. & Brown, P. New insights into the relationship between dopamine, beta oscillations and motor function. Trends in Neurosciences 34, 611–618 (2011).

43. Little, S. & Brown, P. The functional role of beta oscillations in Parkinson’s disease. Parkinsonism & Related Disorders 20, S44–S48 (2014).

44. Tinkhauser, G. et al. The modulatory effect of adaptive deep brain stimulation on beta bursts in Parkinson’s disease. Brain 140, 1053–1067 (2017).

45. Tinkhauser, G. et al. Beta burst dynamics in Parkinson’s disease OFF and ON dopaminergic medication. Brain 140, 2968–2981 (2017).

46. Bonnefond, M. & Jensen, O. Alpha Oscillations Serve to Protect Working Memory Maintenance against Anticipated Distracters. Current Biology 22, 1969–1974 (2012).

47. Jensen, O. & Mazaheri, A. Shaping functional architecture by oscillatory alpha activity: gating by inhibition. Front Hum Neurosci 4, 186 (2010).

48. Rosanova, M. et al. Natural Frequencies of Human Corticothalamic Circuits. J. Neurosci. 29, 7679– 7685 (2009).

49. Fries, P. A mechanism for cognitive dynamics: neuronal communication through neuronal coherence. Trends in Cognitive Sciences 9, 474–480 (2005).

50. Roelfsema, P. R. Solving the binding problem: Assemblies form when neurons enhance their firing rate—they don’t need to oscillate or synchronize. Neuron 111, 1003–1019 (2023).

51. Montemurro, M. A., Rasch, M. J., Murayama, Y., Logothetis, N. K. & Panzeri, S. Phase-of-Firing Coding of Natural Visual Stimuli in Primary Visual Cortex. Current Biology 18, 375–380 (2008).

52. Fries, P., Nikolić, D. & Singer, W. The gamma cycle. Trends in Neurosciences 30, 309–316 (2007).

53. Huang, W. A. et al. Causal oscillations in the visual thalamo-cortical network in sustained attention in ferrets. Current Biology 0, (2024).

54. Pan, Y., Popov, T., Frisson, S. & Jensen, O. Saccades are locked to the phase of alpha oscillations during natural reading. PLOS Biology 21, e3001968 (2023).

55. O’Keefe, J. & Recce, M. L. Phase relationship between hippocampal place units and the EEG theta rhythm. Hippocampus 3, 317–330 (1993).

56. Bosman, C. A. et al. Attentional Stimulus Selection through Selective Synchronization between Monkey Visual Areas. Neuron 75, 875–888 (2012).

57. Grothe, I., Neitzel, S. D., Mandon, S. & Kreiter, A. K. Switching Neuronal Inputs by Differential Modulations of Gamma-Band Phase-Coherence. J. Neurosci. 32, 16172–16180 (2012).

58. Murthy, V. N. & Fetz, E. E. Coherent 25- to 35-Hz oscillations in the sensorimotor cortex of awake behaving monkeys. PNAS 89, 5670–5674 (1992).

59. Howe, M. W., Atallah, H. E., McCool, A., Gibson, D. J. & Graybiel, A. M. Habit learning is associated with major shifts in frequencies of oscillatory activity and synchronized spike firing in striatum. Proceedings of the National Academy of Sciences 108, 16801–16806 (2011).

60. Pfurtscheller, G., Stancák, A. & Neuper, C. Post-movement beta synchronization. A correlate of an idling motor area? Electroencephalography and Clinical Neurophysiology 98, 281–293 (1996).

61. Engel, A. K. & Fries, P. Beta-band oscillations — signalling the status quo? Current Opinion in Neurobiology 20, 156–165 (2010).

62. Jana, S., Hannah, R., Muralidharan, V. & Aron, A. R. Temporal cascade of frontal, motor and muscle processes underlying human action-stopping. eLife 9, e50371 (2020).

63. Pfurtscheller, G., Stancák, A. & Neuper, Ch. Event-related synchronization (ERS) in the alpha band — an electrophysiological correlate of cortical idling: A review. International Journal of Psychophysiology 24, 39–46 (1996).

64. David, O., Kilner, J. M. & Friston, K. J. Mechanisms of evoked and induced responses in MEG/EEG. NeuroImage 31, 1580–1591 (2006).

65. Seedat, Z. A. et al. The role of transient spectral ‘bursts’ in functional connectivity: A magnetoencephalography study. NeuroImage 209, 116537 (2020).

66. Luczak, A., McNaughton, B. L. & Harris, K. D. Packet-based communication in the cortex. Nat Rev Neurosci 16, 745–755 (2015).

67. Leske, S. & Dalal, S. S. Reducing power line noise in EEG and MEG data via spectrum interpolation. NeuroImage 189, 763–776 (2019).

68. Whittington, M. A., Traub, R. D., Kopell, N., Ermentrout, B. & Buhl, E. H. Inhibition-based rhythms: experimental and mathematical observations on network dynamics. International Journal of Psychophysiology 38, 315–336 (2000).

69. Lőrincz, M. L., Kékesi, K. A., Juhász, G., Crunelli, V. & Hughes, S. W. Temporal Framing of Thalamic Relay-Mode Firing by Phasic Inhibition during the Alpha Rhythm. Neuron 63, 683–696 (2009).

70. Wang, X.-J. Neurophysiological and Computational Principles of Cortical Rhythms in Cognition. Physiological Reviews 90, 1195–1268 (2010).

71. Oostenveld, R., Fries, P., Maris, E. & Schoffelen, J.-M. FieldTrip: Open Source Software for Advanced Analysis of MEG, EEG, and Invasive Electrophysiological Data. Intell. Neuroscience 2011, 1:1–1:9 (2011).

72. Shafto, M. A. et al. The Cambridge Centre for Ageing and Neuroscience (Cam-CAN) study protocol: a cross-sectional, lifespan, multidisciplinary examination of healthy cognitive ageing. BMC Neurol 14, 204 (2014).

73. Taulu, S. & Simola, J. Spatiotemporal signal space separation method for rejecting nearby interference in MEG measurements. Phys. Med. Biol. 51, 1759 (2006).

74. Venema, V., Ament, F. & Simmer, C. A Stochastic Iterative Amplitude Adjusted Fourier Transform algorithm with improved accuracy. Nonlinear Processes in Geophysics 13, 321–328 (2006).

75. Theiler, J., Eubank, S., Longtin, A., Galdrikian, B. & Doyne Farmer, J. Testing for nonlinearity in time series: the method of surrogate data. Physica D: Nonlinear Phenomena 58, 77–94 (1992).

76. Schreiber, T. & Schmitz, A. Improved Surrogate Data for Nonlinearity Tests. Phys. Rev. Lett. 77, 635–638 (1996).

77. Venema, V. Surrogate time series and fields. https://ch.mathworks.com/matlabcentral/fileexchange/4783-surrogate-time-series-and-fields (2023).

78. Karvat, G. et al. Real-time detection of neural oscillation bursts allows behaviourally relevant neurofeedback. Commun Biol 3, 1–10 (2020).

79. Apšvalka, D., Ferreira, C. S., Schmitz, T. W., Rowe, J. B. & Anderson, M. C. Dynamic targeting enables domain-general inhibitory control over action and thought by the prefrontal cortex. Nat Commun 13, 274 (2022).

80. Rossi, S., Hallett, M., Rossini, P. M. & Pascual-Leone, A. Safety, ethical considerations, and application guidelines for the use of transcranial magnetic stimulation in clinical practice and research. Clinical Neurophysiology 120, 2008–2039 (2009).

81. Brainard, D. H. The Psychophysics Toolbox. Spatial Vision 10, 433–436 (1997).

82. Rogasch, N. C. et al. Analysing concurrent transcranial magnetic stimulation and electroencephalographic data: A review and introduction to the open-source TESA software. NeuroImage 147, 934–951 (2017).

83. Zmeykina, E., Mittner, M., Paulus, W. & Turi, Z. Weak rTMS-induced electric fields produce neural entrainment in humans. Sci Rep 10, 11994 (2020).

84. Benjamini, Y. & Hochberg, Y. Controlling the False Discovery Rate: A Practical and Powerful Approach to Multiple Testing. Journal of the Royal Statistical Society: Series B (Methodological) 57, 289–300 (1995).

85. Krekelberg, B. BayesFactor: Release 2022 (v2.3.0). Zenodo 10.5281/zenodo.7006300 (2022).

86. van Doorn, J. et al. The JASP guidelines for conducting and reporting a Bayesian analysis. Psychon Bull Rev 28, 813–826 (2021).

87. Maris, E. & Oostenveld, R. Nonparametric statistical testing of EEG- and MEG-data. Journal of Neuroscience Methods 164, 177–190 (2007).

88. Gerber, E. M. permutest. MATLAB Central File Exchange https://www.mathworks.com/matlabcentral/fileexchange/71737-permutest (2023).

