## Supplemental Material for "Universal rhythmic architecture uncovers distinct modes of neural dynamics"

**This file includes:**

Supplementary Text
Supplementary Table S1
Supplementary Figures S1 to S6
Supplementary References

**Supplementary Text**

Parametrical effects on rhythmicity measures

As mentioned in the Lagged Angle Vector Index (LAVI) section of methods, the founding principle underlying the lagged coherence (LC) analysis is that "real" oscillations are repetitive. Therefore, their periodic waveforms should allow for the prediction of future phases based on the phase of the present one. Based on this principle, Fransen and colleagues [1] defined *rhythmicity* as "the consistency of the phase relations between time points that are separated by some interval (lag)". Practically, rhythmicity (computed with LC) measures the predictability of future phases based on the current one: the better future phases can be predicted, the more rhythmic the signal. Fransen et al also noted, that LC varies as a function of both frequency and lag. LC peaks at frequencies corresponding to "canonical bands" (Fig. 1), and decreases with increasing lag (fig. S5A, example from 60 sec recordings from one participant in dataset III).

Based on this dependence of LC on both frequency and lag, several studies [1–3] quantified how oscillatory an activity in a specific band is based on the "lifetime" of the LC. That is, for how long (i.e., for how many lags) does the LC exceed a threshold. However, two difficulties arise when adopting this approach: first, the threshold has to be calculated for each frequency separately, which undermines inter-frequency comparisons [2]. Second, it requires computing the LC for numerous lags (e.g. 0 to 20 cycles in steps of 0.1, summing to 201 lags [2] or 2 to 7 cycles, summing to 51 lags [3]). This makes LC computationally very intensive.

Looking for a computationally fast way to utilize rhythmicity to define oscillatory bands, we investigated how different parameters affect LAVI. First, we noticed that the dynamic range of the rhythmicity is lower in short (<=1 cycle) and long (>=3 cycles) lags compared to lags between 1 and 2 cycles (fig. S5B). This effect is due to a ceiling effect in short lags and a floor effect in long lags. Importantly, the median (across frequencies) is lag-dependent and is highly stable across participants (fig. S5C, 60 sec EEG from N=127 participants, datasets I-IV).

Another parameter that affects the median of LAVI is the duration of the sliding window used to calculate the time-frequency transform. We used Wavelet transform, hence this parameter is reflected in the length of the Wavelet, measured in cycles per frequency. As expected, a wider Wavelet leads to higher assessed rhythmicity (Fig. S5D), since a longer window enforces greater phase consistency by design. This effect reflects an **analytic property** of the wavelet transform rather than a physiological signal. However, our approach does not rely on absolute LAVI values, but rather on **within-subject, within-analysis relative comparisons**, such as deviations from the median at each frequency. Based on the finding that the LAVI median is highly stable across participants (e.g., standard deviation of 0.01, when using lag=1.5 cycles and Wavelet width =5 cycles), and depends on parameters controlled by the experimenter, we developed an approach to assess band borders based on one lag. That is, instead of measuring how rhythmicity changes over time relative to a baseline time-point, we measure how rhythmicity in each frequency changes relative to the median, in one time-point. We found that using a lag of 1.5 cycles and Wavelet of 5 cycles (arrows in fig S5C and S5D) allows estimation of rhythmicity that is computationally efficient, stable across participants, and offers good dynamic range (median of 0.4, that allows frequency to go above and below the median, without ceiling and floor effects).

Importantly, the median of LAVI depends on the lag and wavelet width, regardless of its source. This can be demonstrated with analysis of surrogate (random) data with spectral characteristics similar to the characteristic of neurophysiological data. In general, the power of the electrophysiological signal is inversely proportional to the frequency, or $P = 1/f^{\alpha}$, with P designating Power and the fractal $\alpha$

represent how steeply the power decreases. In brain signals, the fractal (also called the aperiodic component) is predominantly between 0.5 and 2, which is representative for signals dubbed "pink noise" (in the data analysed for this manuscript, out of 1150 recording sites, 82.4% had $0.5 <= \alpha <=2$, fig S5E). Leveraging this feature, we characterized the rhythmicity of data with similar pink-noise characteristics as our recorded data, but with shuffled (that is, random) phases, using the IAAFT algorithm [4–6]. Starting with raw EEG data from participants, we calculated the aperiodic component of the signal by fitting a power function (fig. S5F). Then, we used the fit as the spectrum input to the IAAFT algorithm (fig. S5G). Finally, we computed the rhythmicity (LAVI) of both the original and surrogate data. As shown in fig. S5H, the rhythmicity curves of both "real" and surrogate data share the same median (~0.4, when using lag = 1.5 cycles and Wavelet width = 5 cycles). However, while the "real" data exhibits fluctuations with distinct peaks and troughs, the surrogate ("pink") data fluctuates much less. We further utilized this finding to define noise levels of per-subject LAVI, treating the median as the baseline and matched pink-noise fluctuations as noise-floor. This approach allows statistical inference of each band (see ABBA in the main text, Methods).

Notably, fluctuations in LAVI can be affected by how oscillatory or bursty the data is in specific frequencies, but also from other parameters. These parameters can be exploited do define the noise-floor. First, the higher the aperiodic exponent ($\alpha$), the higher the noise, especially in low frequencies (fig. S5I). Second, noise level decreases (and in fact, accuracy increases) with increased sampling frequency. This effect becomes more prominent with higher frequencies, since the sampling frequency defines how many samples are there in each cycle, and hence the range of possible phases that can be measured (fig. S5J). Finally, the total duration of the recorded session affects noise levels in low frequencies (fig. S5K). Based on this parameterization, one can generate a table containing expected noise levels based on given aperiodic exponent, sampling frequency, and session duration. Generating such table requires simulating at least 20 repetitions of pink-noise for each parameter combination (to allow bootstrapping with $\alpha$=0.05). This process can be time consuming, but carries the advantage of after being done once, the table can be used with all datasets with similar experimental conditions, thus speed up analysis. We make the table created for this manuscript, along with the code used to generate it, publicly available.

### Supplementary Table S1.

| Fig. | Measure | Groups | Statistical test | Test statistic | Deg. of freedom | p-value (corrected) | Effect size | Post-hoc comparisons | Bayes Factor |
|---|---|---|---|---|---|---|---|---|---|
| 3B | Burst duration (power) | Low-High rhythmicity | ANOVA | F = 36.30 | 3,140 | **2.04E-17 (6.1E-17)** | $\eta^2$= 0.437 | Low-high: **9.8E-18**<br>Low-pink: **0.030**<br>High-pink: **1.6E-10**<br>Pink-pink: 0.447 | 3.36E+14 |
| 3C | Burst duration (WTPL) | Low-High rhythmicity | ANOVA | F = 76.01 | 3,140 | **3.11E-29 (1.9E-28)** | $\eta^2$= 0.620 | Low-high: **1.0E-30**<br>Low-pink: **1.3E-08**<br>High-pink: **2.4E-16**<br>Pink-pink: 0.665 | 1.31E+26 |
| 3D | Burst rate | Low-High rhythmicity | ANOVA | F = 33.21 | 3,140 | **2.83E-16 (5.7E-16)** | $\eta^2$= 0.416 | Low-high: **2.7E-17**<br>Low-pink: **3.3E-04**<br>High-pink: **2.0E-10**<br>Pink-pink: 0.390 | 2.58E+13 |
| 3E | Occupancy | Low-High rhythmicity | ANOVA | F = 3.76 | 3,140 | **0.012 (0.015)** | $\eta^2$= 0.075 | Low-high: 0.231<br>Low-pink: 0.400<br>High-pink: **2.6E-03**<br>Pink-pink: **7.6E-03** | 1.401 |
| 3F | Power (relative threshold) | Low-High rhythmicity | ANOVA | F = 18.31 | 3,140 | **4.46E-10 (6.7E-10)** | $\eta^2$= 0.282 | Low-high: **5.0E-03**<br>Low-pink: **4.2E-05**<br>High-pink: **1.9E-06**<br>Pink-pink: **0.036** | 2.4E+12 |
| 3G | Band consistency | Low-High rhythmicity | ANOVA | F = 1.86 | 3,140 | 0.139 (0.139) | $\eta^2$= 0.038 | Low-high: 0.076<br>Low-pink: 0.758<br>High-pink: 0.028<br>Pink-pink: 0.455 | 0.134 |
| 4A | Frequency- $\delta/\theta$ | Dataset | ANOVA | F = 2.50 | 5,172 | 0.033 (0.09) | $\eta^2$= 0.080 | I-IV: **0.043**<br>II-III: **0.021**<br>II-IV: **0.001** | 0.413 |
| 4A | Frequency- $\beta 2$ | Dataset | ANOVA | F = 2.92 | 5,172 | 0.015 (0.059) | $\eta^2$= 0.078 | I-IV: **0.030**<br>I-V: **0.003**<br>III-V: **0.016**<br>V-VI: **0.009** | 0.748 |
| 4A | Frequency- $\gamma 1$ | Dataset | ANOVA | F = 5.12 | 5,172 | **2.1E-04 (0.002)** | $\eta^2$= 0.132 | I-IV: **0.046**<br>I-V: **1.8E-05**<br>II-V: **1.7E-03**<br>III-V: **6.1E-05**<br>IV-V: **0.032**<br>V-VI: **3.2E-04** | 53.664 |
| 4D | Frequency- $\delta/\theta$ | Region | ANOVA | F = 3.52 | 5,313 (only sig.) | **4.1E-03 (0.007)** | $\eta^2$= 0.053 | Occ-VT: **0.003**<br>Occ-Par: **0.001**<br>Occ-PFC: **0.005**<br>Occ-LT: **0.038**<br>Par-SM: **0.021** | 1.271 |
| 4D | Frequency- $\theta/\alpha$ | Region | ANOVA | F = 4.51 | 5, 626 (only sig.) | **4.8E-04 (9.6E-04)** | $\eta^2$= 0.035 | Occ-VT: **9.4E-06**<br>Occ-Par: **0.002**<br>Occ-SM: **0.025**<br>VT-PFC: **0.034**<br>VT-LT: **0.005** | 4.791 |
| 4D | Frequency- $\alpha$ | Region | ANOVA | F = 11.61 | 5, 715 (only sig.) | **8.5E-11 (6.8E-10)** | $\eta^2$= 0.075 | Occ-VT: **4.7E-13**<br>Occ-Par: **4.1E-03**<br>Occ-PFC: **3.9E-04**<br>Occ-SM: **2.0E-05**<br>Occ-LT: **1.8E-05**<br>VT-Par: **1.7E-04**<br>VT-LT: **0.031** | 5.08E+07 |
| 4D | Frequency- $\beta 1$ | Region | ANOVA | F = 5.93 | 5, 641 (only sig.) | **2.3E-05 (6.5E-05)** | $\eta^2$= 0.044 | Occ-VT: **6.3E-04**<br>Occ-PFC: **3.8E-05**<br>Occ-SM: **3.1E-04**<br>Occ-LT: **4.4E-04**<br>VT-Par: **0.025**<br>Par-PFC: **1.7E-03**<br>Par-SM: **0.008** | 100.93 |

| Fig. | Measure | Groups | Statistical test | Test statistic | Deg. of freedom | p-value (corrected) | Effect size | Post-hoc comparisons | Bayes Factor |
|---|---|---|---|---|---|---|---|---|---|
| | | | | | | | | Par-LT: **0.017** | |
| 4D | Frequency-<br>β2 | Region | ANOVA | F = 5.89 | 5, 650<br>(only sig.) | **2.4E-05**<br>**(6.5E-05)** | $\eta^2$= 0.043 | Occ-VT: **1.6E-06**<br>Occ-SM: **0.021**<br>Occ-LT: **0.013**<br>VT-Par: **2.5E-03**<br>VT-PFC: **3.0E-04** | 114.29 |
| 4D | Frequency-<br>γ1 | Region | ANOVA | F = 3.05 | 5, 518<br>(only sig.) | **0.010**<br>**(0.014)** | $\eta^2$= 0.029 | VT-PFC: **0.010**<br>Par-PFC: **5.4E-04**<br>Par-SM: **8.1E-03**<br>PFC-LT: **0.025** | 0.19 |
| 4G | Frequency-<br>δ/θ | Region | ANOVA | F = 2.50 | 6,501<br>(only sig.) | **0.022**<br>**(0.034)** | $\eta^2$= 0.029 | MF-LT: **0.024**<br>LF-LT: **0.009**<br>Ins.-LT: **0.006**<br>Ins.-MT: **0.033** | 0.11 |
| 4G | Frequency-<br>α | Region | ANOVA | F = 5.78 | 6, 726<br>(only sig.) | **6.8E-06**<br>**(1.0E-04)** | $\eta^2$= 0.046 | MF-LF: **0.004**<br>LF-Ins.: **0.004**<br>LF-MP: **0.023**<br>LF-LP: **1.8E-06**<br>LF-LT: **3.5E-06**<br>LF-MT: **1.0E-06** | 439.94 |
| 4G | Frequency-<br>β1 | Region | ANOVA | F = 5.41 | 6, 726<br>(only sig.) | **1.7E-05**<br>**(1.0E-04)** | $\eta^2$= 0.043 | MF-LF: **8.5E-04**<br>LF-Ins.: **0.025**<br>LF-MP: **0.010**<br>LF-LP: **1.5E-06**<br>LF-LT: **5.6E-06**<br>LF-MT: **2.0E-05** | 159.91 |
| 4G | Frequency-<br>β2 | Region | ANOVA | F = 3.93 | 6, 721<br>(only sig.) | **7.2E-04**<br>**(1.9E-03)** | $\eta^2$= 0.032 | MF-LF: **4.3E-02**<br>LF-LP: **4.7E-05**<br>MP-LP: **0.024**<br>LP-LT: **5.5E-04**<br>LP-MT: **1.6E-02** | 2.94 |
| 4G | Frequency-<br>γ1 | Region | ANOVA | F = 3.28 | 6, 707<br>(only sig.) | **3.4E-03**<br>**(0.007)** | $\eta^2$= 0.027 | MF-LF: **5.0E-02**<br>MF-LT: **2.1E-03**<br>Ins.-LT: **0.010**<br>MP-LT: **8.1E-03**<br>LP-LT: **4.3E-04**<br>LT-MT: **0.023** | 0.50 |
| 5B | Band frequency | Invasive – non-invasive | ANOVA-method | F = 0.3 | 1,5527 | 0.58 | $\eta^2$= 5.45E-5 | | 0.042 |
| 5D | Band frequency | Rat – Human | ANOVA-species | F = 0.05 | 1,336 | 0.83 | $\eta^2$=1.43E-4 | | 0.091 |
| 5F | Band frequency | Female - Male | ANOVA-sex | 0.26 | 1,4414 | 0.611 | $\eta^2$=5.9E-05 | | 0.0231 |
| 6B | Band frequency | Age | Pearson correlation | ρ =<br>δ: –0.149<br>δ/θ: –0.040<br>θ: –0.208<br>θ/α: –0.228<br>α: –0.340<br>β1: –0.119<br>β2: –0.134<br>γ1: 0.015 | n =<br>δ: 253<br>δ/θ: 539<br>θ: 590<br>θ/α: 618<br>α: 618<br>β1: 618<br>β2: 612<br>γ1: 612 | corrected<br>**δ: 0.024**<br>δ/θ: 0.409<br>**θ: 9.8E–07**<br>**θ/α: 4E–08**<br>**α: 2.9E–17**<br>**β1: 5E–03**<br>**β2: 2E–03**<br>γ1: 0.715 | | | |
| 6B | Band rhythmicity magnitude | Age | Pearson correlation | ρ =<br>δ: 0.147<br>δ/θ: –0.058<br>θ: –0.051<br>θ/α: 0.057<br>α: –0.203<br>β1: 0.333<br>β2: –0.029<br>γ1: –0.012 | n =<br>δ: 253<br>δ/θ: 539<br>θ: 590<br>θ/α: 618<br>α: 618<br>β1: 618<br>β2: 612<br>γ1: 612 | corrected<br>δ: 0.052<br>δ/θ: 0.283<br>θ: 0.286<br>θ/α: 0.283<br>**α: 1.5E–06**<br>**β1:1.6E–16**<br>β2: 0.543<br>γ1: 0.770 | | | |
| 6B | Band frequency | Age | ANOVA-<br>δ | F = 1.95 | 6,244 | 0.074<br>(0.099) | $\eta^2$= 0.046 | | 0.066 |

| Fig. | Measure | Groups | Statistical test | Test statistic | Deg. of freedom | p-value (corrected) | Effect size | Post-hoc comparisons | Bayes Factor |
|---|---|---|---|---|---|---|---|---|---|
| 6B | Band frequency | Age | ANOVA-$\delta/\theta$ | F = 0.73 | 6,525 | 0.623 (0.712) | $\eta^2$= 0.0083 | | 5.37E-4 |
| 6B | Band frequency | Age | ANOVA-$\theta$ | F = 5.1 | 6,574 | **4E-5 (1.07E-4)** | $\eta^2$= 0.05 | 18-28 - 68-78: **3.4E-04**<br>18-28 - 78-88: **8.1E-03**<br>28-38 - 48-58: **0.031**<br>28-38 - 58-68: **0.012**<br>28-38 - 68-78: **5.0E-06**<br>28-38 - 78-88: **6.8E-04**<br>38-48 - 68-78: **5.6E-04**<br>38-48 - 78-88: **0.022**<br>48-58 - 68-78: **0.016**<br>58-68 - 68-78: **0.046** | 55.3 |
| 6B | Band frequency | Age | ANOVA-$\theta/\alpha$ | F = 6.8 | 6,604 | **5.34E-7 (2.14E-6)** | $\eta^2$= 0.063 | 18-28 - 48-58: **0.044**<br>18-28 - 58-68: **0.031**<br>18-28 - 68-78: **1.1E-03**<br>18-28 - 78-88: **7.3E-04**<br>28-38 - 48-58: **7.8E-03**<br>28-38 - 58-68: **4.7E-03**<br>28-38 - 68-78: **4.1E-05**<br>28-38 - 78-88: **2.9E-05**<br>38-48 - 48-58: **8.1E-03**<br>38-48 - 58-68: **4.7E-03**<br>38-48 - 68-78: **3.2E-05**<br>38-48 - 78-88: **2.4E-05** | 4.78E+3 |
| 6B | Band frequency | Age | ANOVA-$\alpha$ | F = 13.73 | 6,604 | **1.18E-14 (9.45e-14)** | $\eta^2$= 0.12 | 18-28 - 48-58: **6.5E-04**<br>18-28 - 58-68: **3.5E-06**<br>18-28 - 68-78: **4.2E-08**<br>18-28 - 78-88: **7.2E-08**<br>28-38 - 48-58: **6.9E-04**<br>28-38 - 58-68: **1.3E-06**<br>28-38 - 68-78: **5.9E-09**<br>28-38 - 78-88: **1.6E-08**<br>38-48 - 48-58: **0.010**<br>38-48 - 58-68: **4.4E-05**<br>38-48 - 68-78: **3.0E-07**<br>38-48 - 78-88: **6.6E-07**<br>48-58 - 68-78: **0.012**<br>48-58 - 78-88: **0.012** | 2.55E+11 |
| 6B | Band frequency | Age | ANOVA-$\beta 1$ | F = 2.73 | 6,604 | **0.0125 (0.025)** | $\eta^2$= 0.0264 | 18-28 - 48-58: **0.047**<br>18-28 - 58-68: **0.020**<br>18-28 - 68-78: **7.4E-03**<br>18-28 - 78-88: **7.3E-03**<br>28-38 - 68-78: **0.020**<br>28-38 - 78-88: **0.020** | 0.082 |
| 6B | Band frequency | Age | ANOVA-$\beta 2$ | F = 2.27 | 6,598 | 0.035 (0.057) | $\eta^2$= 0.0223 | | 0.0223 |
| 6B | Band frequency | Age | ANOVA-$\gamma 1$ | F = 0.56 | 6,598 | 0.762 (0.762) | $\eta^2$= 0.0056 | | 2.23E-4 |
| 6D | Burst rate | PD, Bands x On/Off med | ANOVA2-bands | F = 23.78 | 1,60 | **8.3E-6** | $\eta^2$= 0.28 | $\beta$1,off–$\beta$2,off: **2.0E-03**<br>$\beta$1,off–$\beta$2,on: **1.6E-03**<br>$\beta$1,on–$\beta$2,off: **0.018**<br>$\beta$1,off–$\beta$2,on: **0.015** | 2.83E+3 |
| 6D | Burst rate | PD, Bands x On/Off med | ANOVA2-medication | F = 0.33 | 1,60 | 0.56 | $\eta^2$= 0.056 | | 0.18 |
| 6D | Burst rate | PD, Bands x On/Off med | ANOVA-2 interaction | F = 0.23 | 1,60 | 0.64 | $\eta^2$= 0.004 | | 0.246 |
| 6E | Burst duration | PD, Bands x On/Off med | ANOVA2-bands | F = 0.51 | 1,60 | 0.48 | $\eta^2$= 0.009 | | 0.204 |
| 6E | Burst duration | PD, Bands x On/Off med | ANOVA2-medication | F = 10.59 | 1,60 | **0.0019** | $\eta^2$= 0.154 | $\beta$1,off–$\beta$1,on: **0.0026**<br>$\beta$1,off–$\beta$2,on: **0.007** | 14.875 |
| 6E | Burst duration | PD, Bands x On/Off med | ANOVA2-interaction | F = 1.605 | 1,60 | 0.21 | $\eta^2$= 0.027 | | 0.456 |

**Statistical reporting table.**

All tests are two-sided. Correction for multiple comparisons was performed with the false discovery rate according to the Benjamini-Hochberg algorithm. Post-hoc tests following ANOVA were performed with the Fisher's least significant difference procedure. Significant p values are presented in bold. Fig.: the figure panel in which the test is presented. Deg.: degrees.

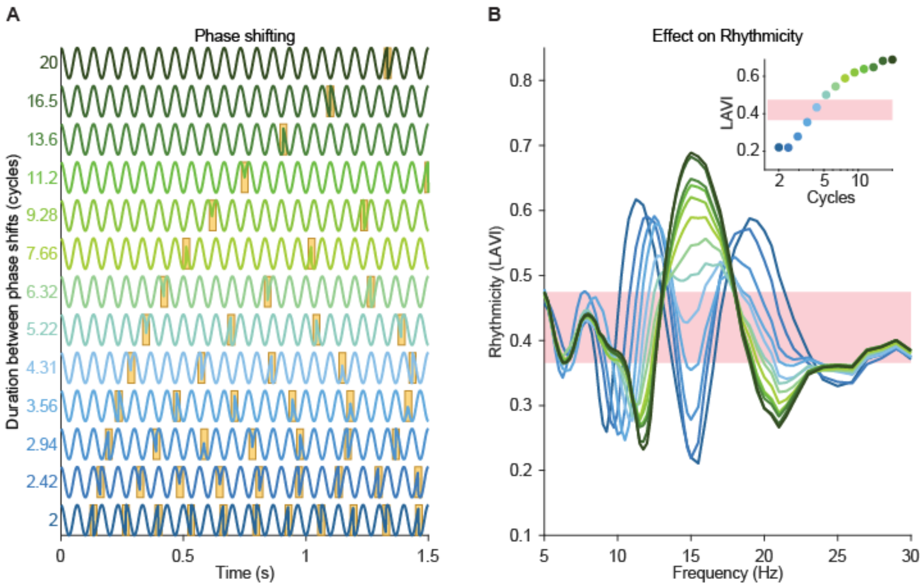


**Supplementary Figure S1. The Lagged Angle Vector Index (LAVI) measures rhythmicity.**

In each frequency, LAVI reflects how sustained the oscillation is over the session: sustained oscillations receive high rhythmicity values, while frequent phase shifts reduce LAVI. For direct demonstration of the effect of sustainability on LAVI, we flipped the sign of the phase of the 15 Hz component of pink-noise every set amount of cycles ranging from 2 to 20. (**A**) Simulated signal, filtered at 15 Hz, arranged from "ephemeral" (blue) to "rhythmic" (green). Phase-shifts are marked in gold. (**B**) The LAVI profile over frequencies. Colour of traces corresponds to (**A**). Pink shading corresponds to the noise, estimated as the minimal and maximal LAVI values of the pink-noise data (without filter weight adjustment at 15 Hz). Inset: LAVI values at the modulated frequency (15 Hz).The signal becomes more rhythmic than noise around four cycles. Note that increasing (decreasing) the rhythmicity in one band decreases (increases) rhythmicity in neighbouring bands. This can be caused by interference. However, peaks and troughs in physiological data are above and beyond this effect (for more details, see fig. S3).

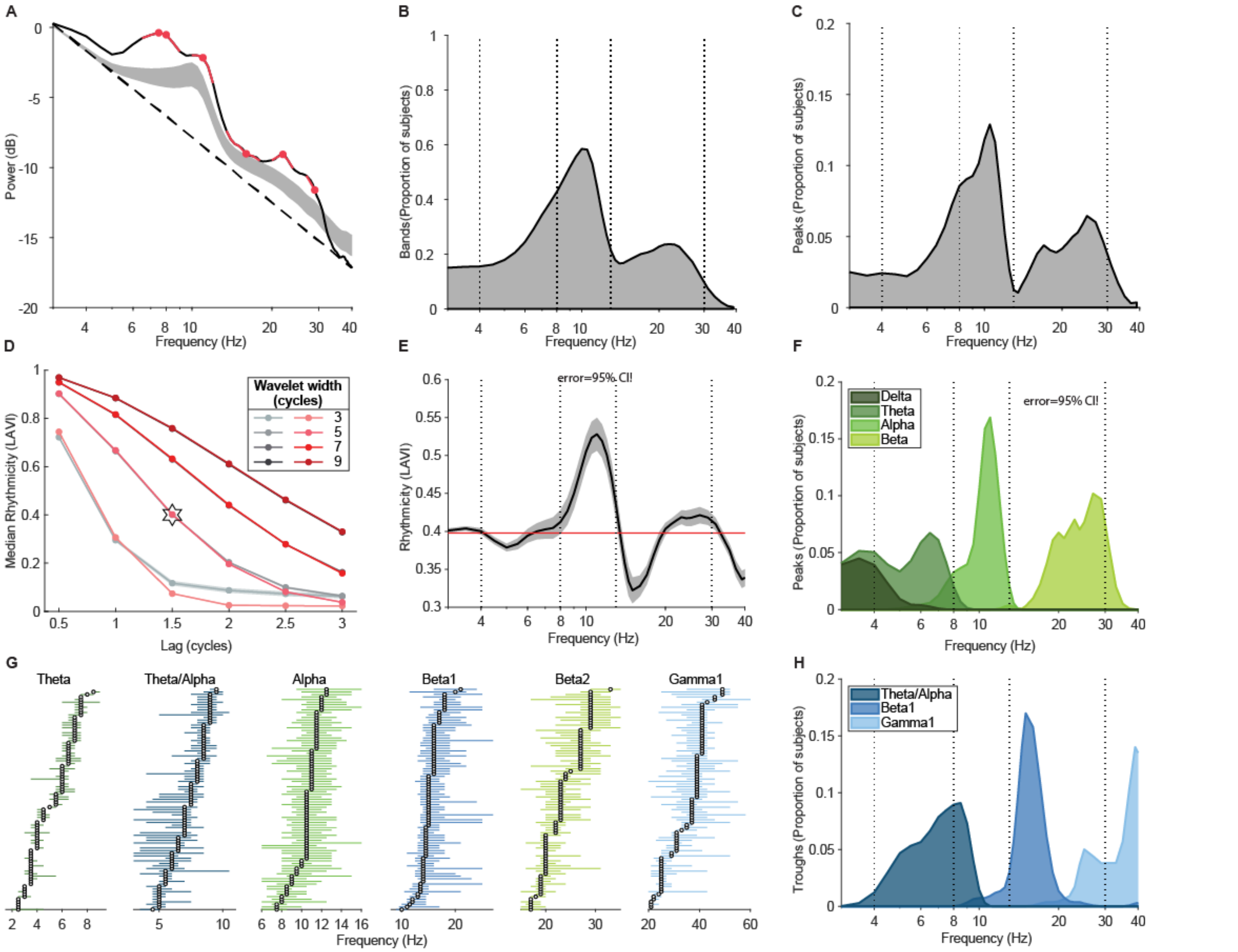


**Supplementary Figure S2. Band segregation using power and rhythmicity.**

(**A**) Brainwave bands are commonly defined as bumps above the 1/*f* aperiodic component. Black trace: power spectrum of one subject (same as in Fig. 1C-D). Dashed line: the 1/*f* fit. Pink: bands detected using an iterative Gaussian-fit algorithm (specparam, formerly fooof, ref. S18). Grey shading: 95% confidence-interval (CI) of population power (N = 90 participants from datasets I, II, and IV). Group averaged of bands detected by specparam are presented in (**B**), and peaks in (**C**). Dashed lines in (**B**), (**C**), (**E**), (**F**), and (**H**) at 4,8,12, and 30 Hz denote borders between canonical bands (ref. 17). (**D**) Median rhythmicity levels are stable across participants and data types. Grey (pink) traces denote the mean and shades the 95% CI of the median of 90 subjects (pink-noise) using different lags and wavelet widths. Note that with 5 or more wavelet length, and 2 or less cycle lags, noise and data are indistinguishable. This allows using surrogate data to estimate noise floor and significance levels, as further discussed in Supplementary Text, Parametrical effects on rhythmicity measures, and fig. S4. Star denotes the values used throughout the manuscript. (**E**) Black: Mean ± 95% CI of the population (N = 90) rhythmicity values. Pink: mean ± 95% CI of the medians of each subject. (**F**) Population summary of significantly ($p$<0.05) sustained peaks detected by ABBA. (**G**) Forest plot of significant bands. Circle: peak (or trough) of each subject. Green horizontal lines: limits of rhythmic bands. Blue horizontal lines: limits of arrhythmic bands. Note that although the ranges of bands are stable and agree with canonical bands (ref 17), there is a considerable degree of inter-subject variability, which can lead to erroneous band-definition and deems individual band definition necessary. (**H**). Population summary of significantly ($p$<0.05) transient peak-frequencies (i.e., troughs in LAVI) detected by ABBA.

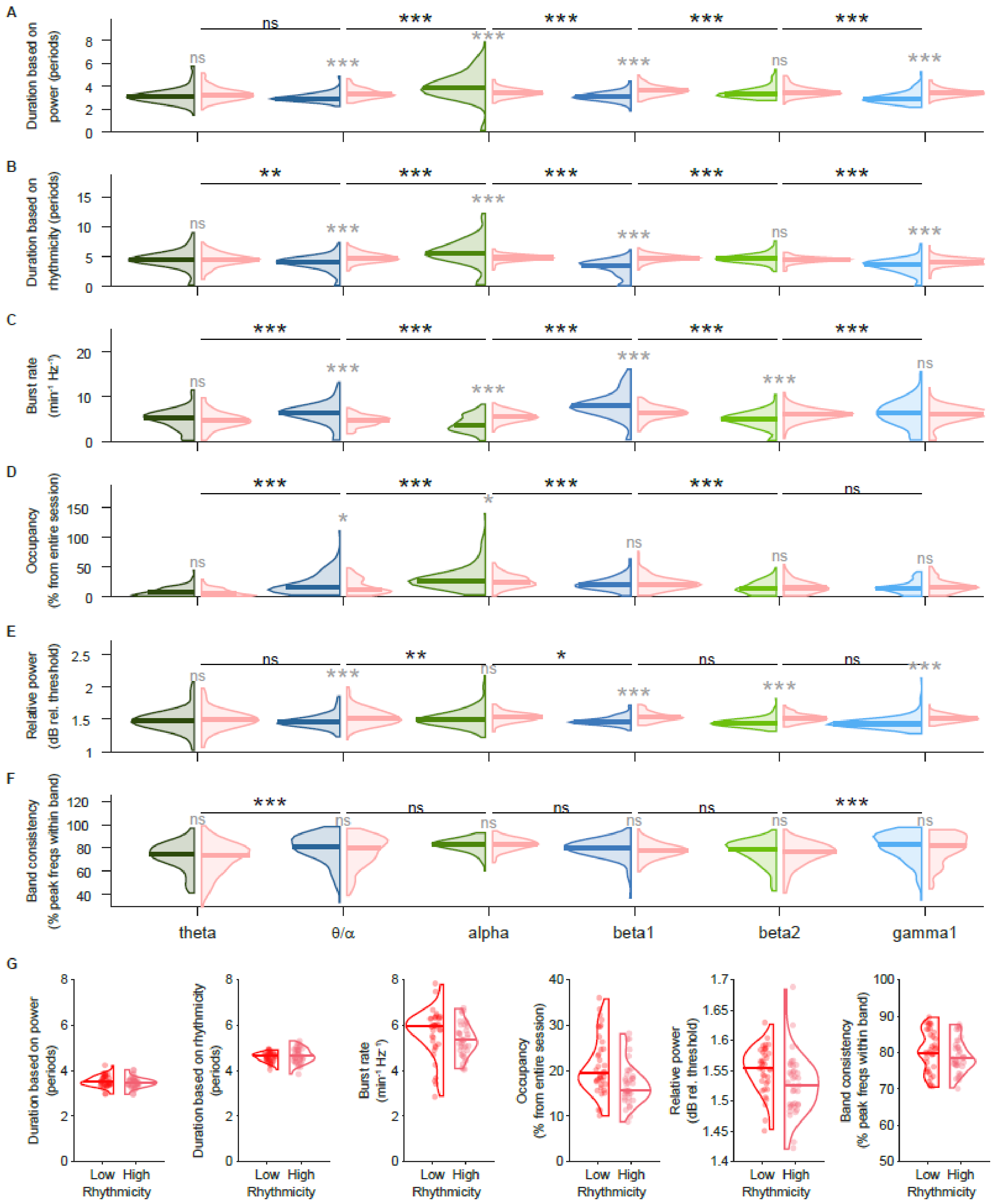


### Supplementary Figure S3. Band-specific burst-statistics.

The duration (**A,B**), rate (**C**), occupancy (**D**), relative power (**E**) and band frequency consistency (**F**) of oscillatory bursts in EEG data (N=178 participants, blue and green distributions, corresponding to bursty and sustained bands respectively) and simulations with participant-matched aperiodic 1/f noise (pink distributions). Violin plots represent the whole distribution, horizontal lines the median. *- p<0.05, **- p<0.01, ***- p<0.001, ns- not significant, ANOVA with Tukey's post-hoc. (**G**)

Comparison of distributions of pink-noise simulations collapsed for low- and high-rhythmicity, from dataset I (compare to recorded data in main Fig.
3B-F).

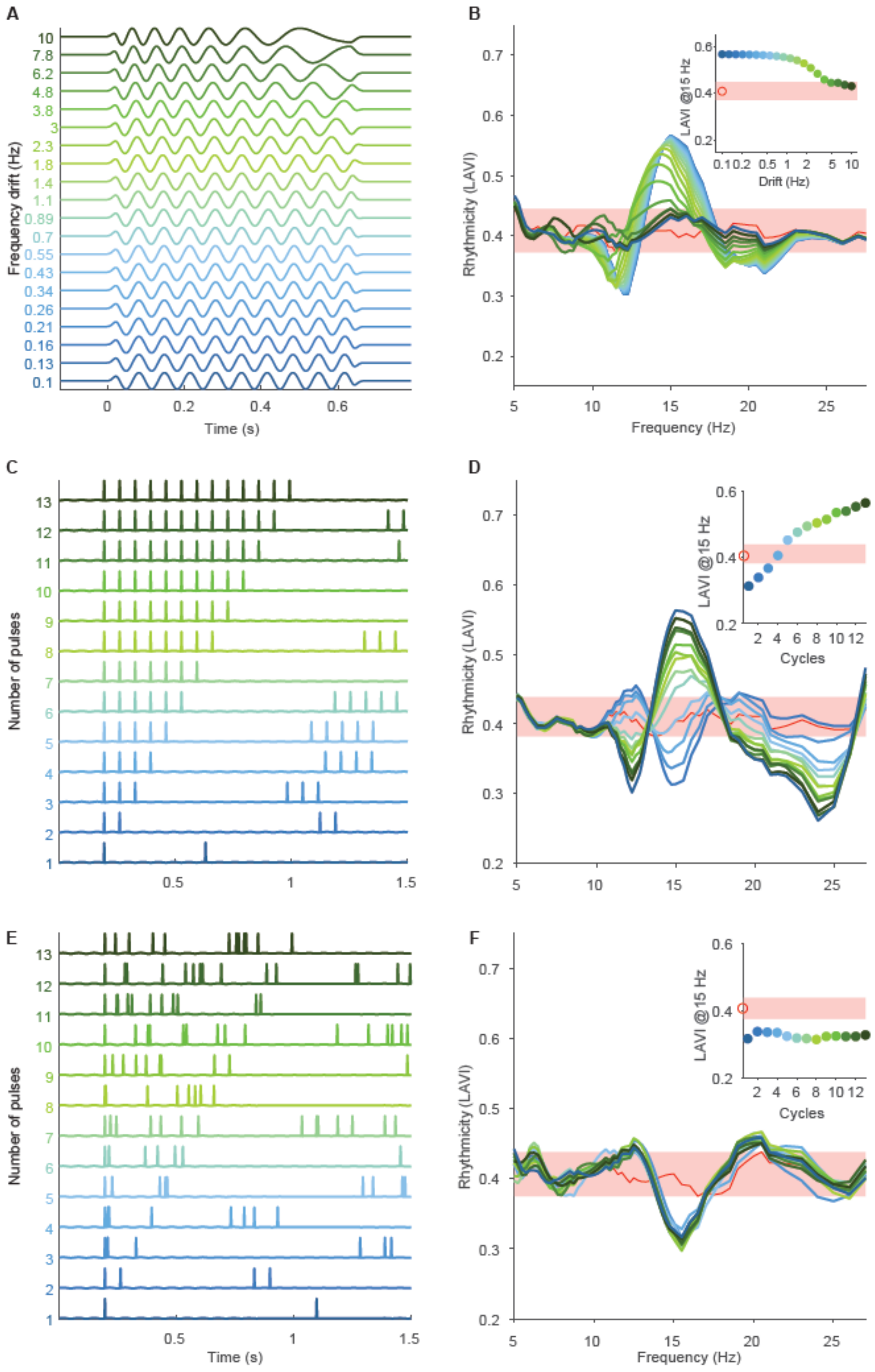


### Supplementary Figure S4. The effect of repeating pulses and frequency drifts on rhythmicity

**A-B**: Bursts with frequency drifts. Each burst consisted of 10 cycles, and the frequency drifted linearly from 15 Hz plus half the value on the y-axis to 15 Hz minus half the value. **B**: the effect on rhythmicity. Note that frequency drift reduces rhythmicity, however, rhythmicity remains above noise-level up to drifts of 5 Hz (33% of original frequency).

**C-D**: Rhythmic pulses, **E-F**: Arrhythmic pulses. **C,E**: Simulations design. Synchronized (**C**) or asynchronized (**E**) transient inputs were simulated as trains of strong pulses. The first and last inputs in a train were time locked to the same duration, whereas the intervals between inputs within trains were constant in **C** and randomized in **E**. **D,F**: The response of the rhythmicity profile to synchronized (**D**) or asynchronized (**F**) inputs. Colour code identical to main Fig. 3H. Note rhtyhmicity levels higher than expected by noise with 5 or more synchronized pulses (**D**), but no effect of number of pulses on rhythmicity with asynchronized pulses (**F**).

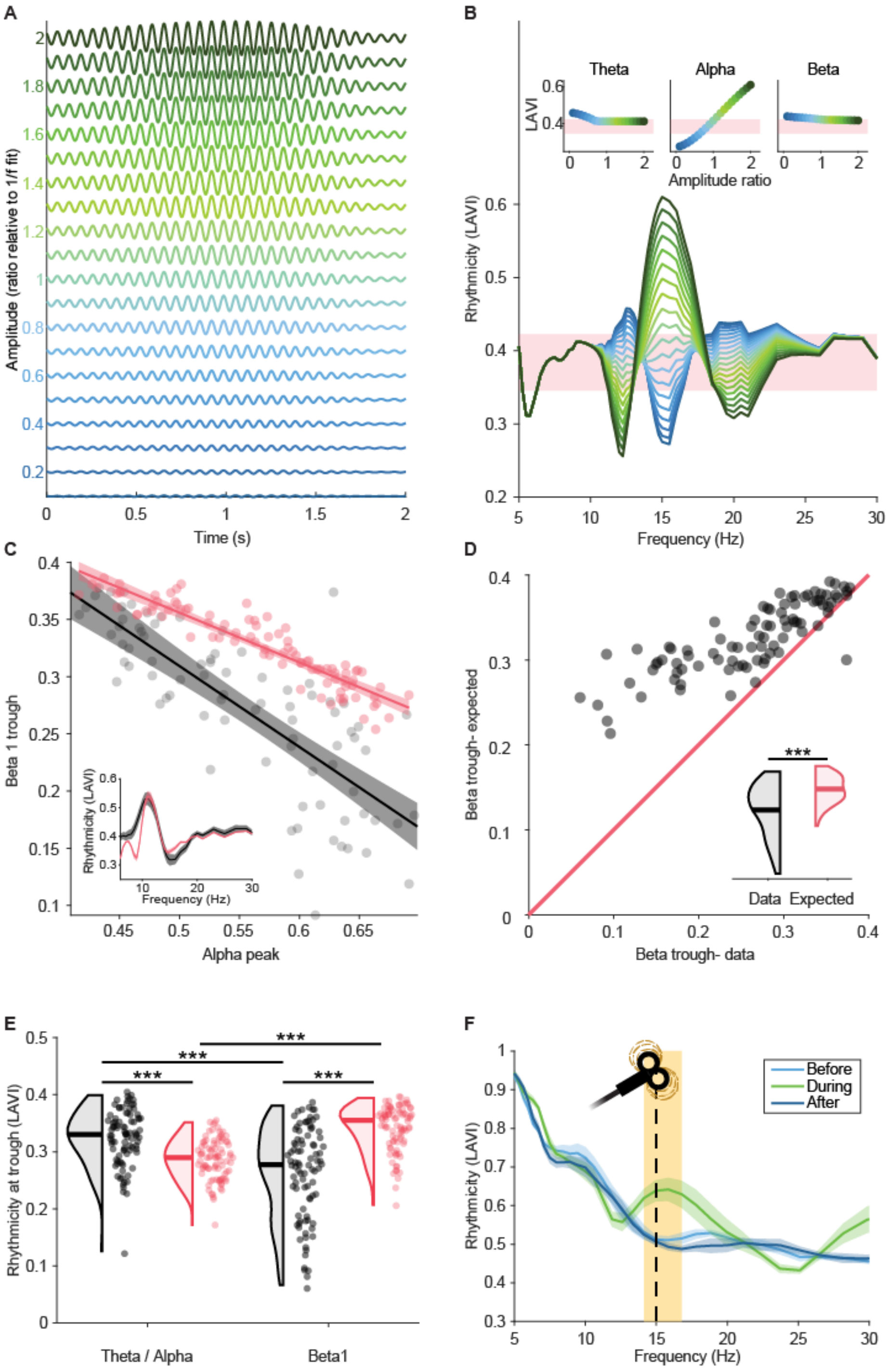


**Supplementary Figure S5. Transient bands are more arrhythmic than expected by interference shadows from neighbouring rhythmic bands.**

(**A-B**) Potential cross-frequency effects on the rhythmicity profile. Reducing the power in one frequency, e.g. by filtering (**A**, blue traces) reduces the rhythmicity in this frequency below expected by pink-noise (**B**), but also increases rhythmicity in neighbouring frequencies. Conversely, increasing the power (green traces) increases rhythmicity in the manipulated frequency and reduces rhythmicity in the neighbouring frequencies. This interference shadow can theoretically be the (artefactual) source of troughs in the rhythmicity profile that would be erroneously defined as arrhythmic bands. (**C**) Rhythmicity levels (LAVI) at the alpha peak vs. the beta1 trough of 90 participants (black) or pink-noise (pink). To generate the pink noise distribution, we created 90 instantiations of 120 s pink noise, each with power at 11 Hz (alpha) randomly chosen between 1.05 to 2.25 of the original (1/*f*) value. Dots: individual subjects/ noise instantiations. Lines: linear regression. Shades: 95% prediction intervals. Data: Pearson $\rho=-0.832$, $p<10^{-23}$. Noise: $\rho=-0.927$, $p<10^{-38}$. Inlet: Population mean ± 95% CI of rhythmicity values. (**D**) Based on the alpha-to-beta1 linear regression of pink-noise, we compared the beta1 rhythmicity levels measured in the data (abscissa) to the beta1 levels expected given the alpha peak levels of each participant (ordinate). Note that 89 out of 90 subjects (99%) were above the unity line, indicating that their beta1 trough is lower than expected by interference shadows from alpha ($t_{89}=11.8$, $p<10^{-19}$, inlet). (**E**) Beta1 trough is more arrhythmic than expected by alpha interference. For each subject we created a 3 min surrogate pink-noise trace, with matched aperiodic component and alpha power (10 Hz) matching the original (non-fit) value, and extracted the rhythmicity levels at the trough between theta and alpha and the beta1 trough of both data (black) and surrogate (pink). Rainclouds show the full distribution. Horizontal lines: median. Two-way ANOVA (band and data type): main effect for band: $F_{1,356}=0.94$, $p=0.33$; main effect for data type: $F_{1,356}=14.1$, $p<10^{-3}$; interaction: $F_{1,356}=95$, $p<10^{-19}$. ***- p<0.001, post-hoc t-test with Bonferroni correction. The significant interaction indicates that the artefactual interference effect of rhythmic alpha is expected to be stronger on lower frequencies, but in the data the beta1 trough is more arrhythmic than expected by noise and in comparison to the theta-alpha trough. (**F**) Repetitive TMS increases rhythmicity in the stimulated frequency, but the interference shadow does not reach significance. The rhythmicity levels before (light blue), during (green) and after (dark blue) 6 rhythmic TMS stimulations at 15 Hz (dashed line). Shaded area: rhythmicity during is different than before/ after ($p<0.05$, cluster permutation test).

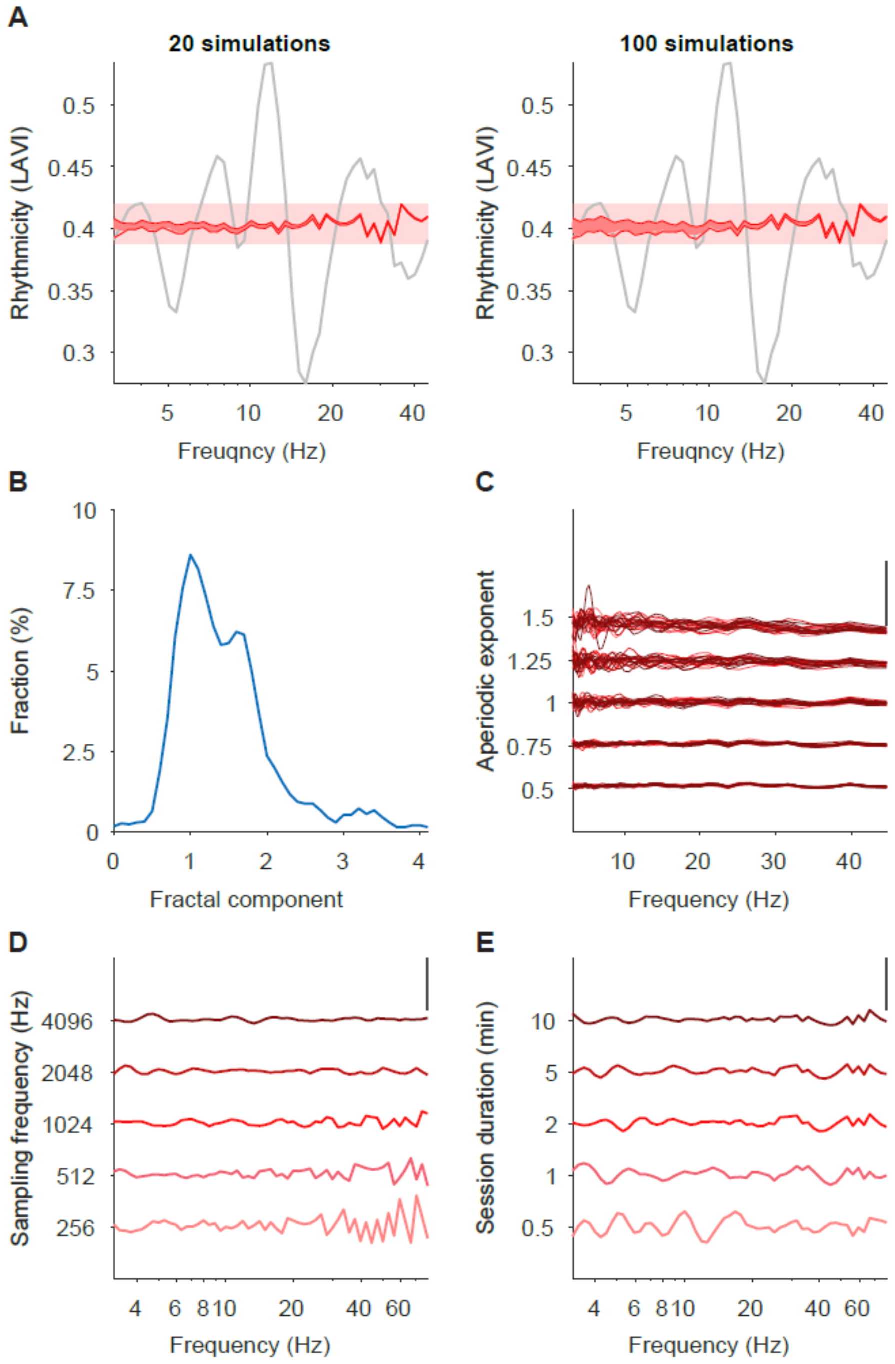


**Supplementary Figure S6. Parameters affecting pink-noise simulation.**

**A**. The number of pink-noise simulations (pink traces) has minimal impact on the estimated noise ribbon. Two approaches are suggested for defining the noise ribbon: (1) per-frequency limits (red traces), and (2) global minimum–maximum values across the spectrum (pink shading). Grey trace shows the LAVI profile of empirical data. In this example (sampling rate = 512 Hz; simulation duration = 3 min; aperiodic exponent = 1.1), the number of simulations slightly affects low frequencies. The second (global) approach is more sensitive to high-frequency variance, but remains stable across simulation counts. Notably, low-frequency variation is primarily driven by the aperiodic exponent (panel C) and session duration (panel E), while high-frequency variability is mainly influenced by the sampling rate (panel D). **B**. Distribution of aperiodic (fractal) exponents of empirical data, with most values falling between 0.5 and 2—typical of "pink" noise. **C-E**. Effects of simulation parameters on the LAVI spectrum.

**C**. Varying the aperiodic exponent.

**D**. Varying the sampling frequency.

**E**. Varying the duration of the simulated session. Verticaltal bars in (C-E) denote LAVI = 0.1.

**Supplementary references**


1. Fransen, A. M. M., van Ede, F. & Maris, E. Identifying neuronal oscillations using rhythmicity. *NeuroImage* **118**, 256–267 (2015).
2. Myrov, V. *et al.* Rhythmicity of neuronal oscillations delineates their cortical and spectral architecture. *Commun Biol* **7**, 1–18 (2024).
3. Rayson, H. *et al.* Detection and analysis of cortical beta bursts in developmental EEG data. *Developmental Cognitive Neuroscience* **54**, 101069 (2022).
4. Venema, V., Ament, F. & Simmer, C. A Stochastic Iterative Amplitude Adjusted Fourier Transform algorithm with improved accuracy. *Nonlinear Processes in Geophysics* **13**, 321–328 (2006).
5. Theiler, J., Eubank, S., Longtin, A., Galdrikian, B. & Doyne Farmer, J. Testing for nonlinearity in time series: the method of surrogate data. *Physica D: Nonlinear Phenomena* **58**, 77–94 (1992).
6. Schreiber, T. & Schmitz, A. Improved Surrogate Data for Nonlinearity Tests. *Phys. Rev. Lett.* **77**, 635–638 (1996).
